# A cortical code emerges in Layer5a of S1 through temporal integration of thalamic inputs

**DOI:** 10.1101/2025.06.11.658272

**Authors:** Tom Quétu, Daniel E. Shulz, Matías A. Goldin

**Affiliations:** Université Paris-Saclay, CNRS, Institut des Neurosciences Paris-Saclay (NeuroPSI), 91400 Saclay, France

## Abstract

Tactile representations in the barrel field of the primary somatosensory cortex (wS1) receive input from two thalamic nuclei: ventro-posterior-medial (VPM) and posterior-medial complex (POm). However, how these inputs generate distinct cortical representations remains unclear. Previous work revealed a sweep-stick code in rat wS1 using novel velocity-white noise whisker stimulus. Sticks are high-velocity single-whisker bumps; sweeps are large multi-whisker displacements with extended temporal profiles. We hypothesized that cortex neurons inherit “stick” responses from VPM and “sweep” responses from first order POm. We tested this by studying coding strategies in both thalamic nuclei and wS1 in mice, determining where the cortical sweep and stick codes originate. We found a stratified wS1 representation of sweep and stick classes, whereas in our recordings VPM and POm mostly contained stick-responding neurons. Layer 4 “stick” responses are delayed versions of VPM responses, and layer 5b “sweep” responses come from first order POm. Notably, layer 5a “sweep” responses arise from temporal integration of “stick” information from VPM and first order POm. Thus, contrary to our initial hypothesis, sweep responses are not inherited from the VPM or first order POm but are constructed within the cortex. In contrast, higher order POm presented only “sweep” responses inherited from layer 5b of cortex, with a latency similar to that in layer 5a. Our findings reveal a circuit where fast VPM-encoded “sticks” allow fine-tuned texture processing, complemented by cortically constructed sweep responses that integrate multi-whisker information.

## Introduction

Recent work on the whisker-to-barrel cortex system has highlighted the central role of the thalamus in both sensory processing and perceptual decision making. It has been shown that even in the absence of the whisker sensory cortex, mice can still detect objects using their whiskers (1), a result that mirrors findings in the auditory system for sound detection (2). These studies suggest that the thalamus alone can support simple detection tasks. Beyond simple detection, the thalamus can flexibly encode sensory input based on experience or context, for instance, differentiating touch and visual inputs depending on conditioning (3). More broadly, thalamocortical loops have been proposed to support complex sensorimotor interactions, such as active sensing (4), highlighting the dynamic nature of thalamic contributions to cortical processing. Across modalities, the thalamus obeys similar connectivity patterns. In the visual system, the thalamic reticular nucleus is connected to bottom-up and top-down pathways with the cortex. This nucleus is involved in the transfer of sensory and motor information as well as attention between the cortex and the thalamus (5). In the whisker system, the POm nucleus appears to have the same role (6).

While many aspects of thalamic responses to whisker input have been characterized, the specific movement features that drive selective responses in their neurons remain not fully understood. The thalamus is structured in multiple regions that involve different functions: VPM primarily encodes single whisker movements and POm encodes multiple whisker input (7). POm itself can be further divided into first-order and higher-order subregions (8, 9). These subdivisions connect to different cortical layers, suggesting distinct functions in sensorimotor processing (10–12).

Diverse coding strategies are already implemented in the thalamus. During a whisker discrimination task, the activity during licking was stronger in POm, implying a role in decision making for this structure (11). Other strategies are believed to take part in VPM to allow encoding precise information about texture and vibrissae dynamics with precise spike timings (13). However, it still remains poorly understood what are the precise stimulus features (that is the precise whisker deflections) encoded in the thalamus, and how they give rise to the sensory features encoded across the cortical layers.

Following this line of inquiry, we studied here the precise characteristics of whisker deflections encoded in both the cortex and the thalamus. This was previously challenging due to two significant obstacles that had to be overcome. First, it is hard to control precisely the movement of the whiskers. Poor control stimulation techniques, such as air puffs impacting multiple whiskers without control, or just a single whisker deflection were used broadly. Second, the definition of the parametric space for stimulus design affects the observed coding properties of cortical neurons. The first problem was solved by the use of a multi-whisker stimulation device, based on piezoelectric benders, to control up to 24 whiskers simultaneously and independently with millisecond and micrometer precision (14). The second problem was overcome with the development of Gaussian white noise simulations in the velocity domain (15). This optimized stimulus elicited the most feature-selective neuronal responses in wS1. These technical advancements previously allowed us to identify, in the rat barrel cortex, a novel coding strategy (15): fast single whisker velocity events called ‘sticks’, and broad multi-whisker high velocity events, called ‘sweeps’, with some evidence that these coding classes are segregated across different cortical layers. That study, however, was restricted to the cortex and did not address where these codes originate. Here we move to the mouse and ask whether the stick and sweep codes are inherited from the thalamus.

To study the origin of ‘stick-sweep’ coding, we recorded multiple single-neuron activity in the barrel cortex and the two thalamic nuclei of mice during optimized multi whisker deflections. Using silicon probes, we first made a thorough study of the responses of all cortical layers to a wide repertoire of stimulations including: rostro-caudal and ventro-dorsal velocity simulations, together with position and acceleration optimized Gaussian white noise. Then, we recorded VPM and first order POm responses to the same set of whisker deflections to explore the potential thalamic inheritance of the cortical coding.

## Results

### A rostro-caudal velocity-optimized stimulus is suited to study wS1 responses

Exhaustively exploring the full parameter space of sensory inputs is often impractical due to the high dimensionality of the stimulus and the limited duration of viable recordings. Instead, incorporating a priori knowledge about neuronal response properties can help target specific stimulus dimensions, thereby revealing fine-tuned coding preferences more efficiently. Building on previous results from our laboratory in rat wS1 (15), we designed optimized stimuli to explore a 2-dimensional space of multi-whisker movements (involving up to 24 whiskers) position (POS_W_), velocity (VEL_W_), and acceleration (ACC_W_) (Fig. 1B) in lightly anesthetized mice (see Methods), while recording neuronal responses in wS1 using multielectrode silicon probes (Fig. 1A, Supplementary Fig. S1A-B). Our stimulus design includes white noise broadband deflections in whisker position (POS_W_), velocity (VEL_W_) and acceleration (ACC_W_) (Fig. 1B).

**Figure 1.**
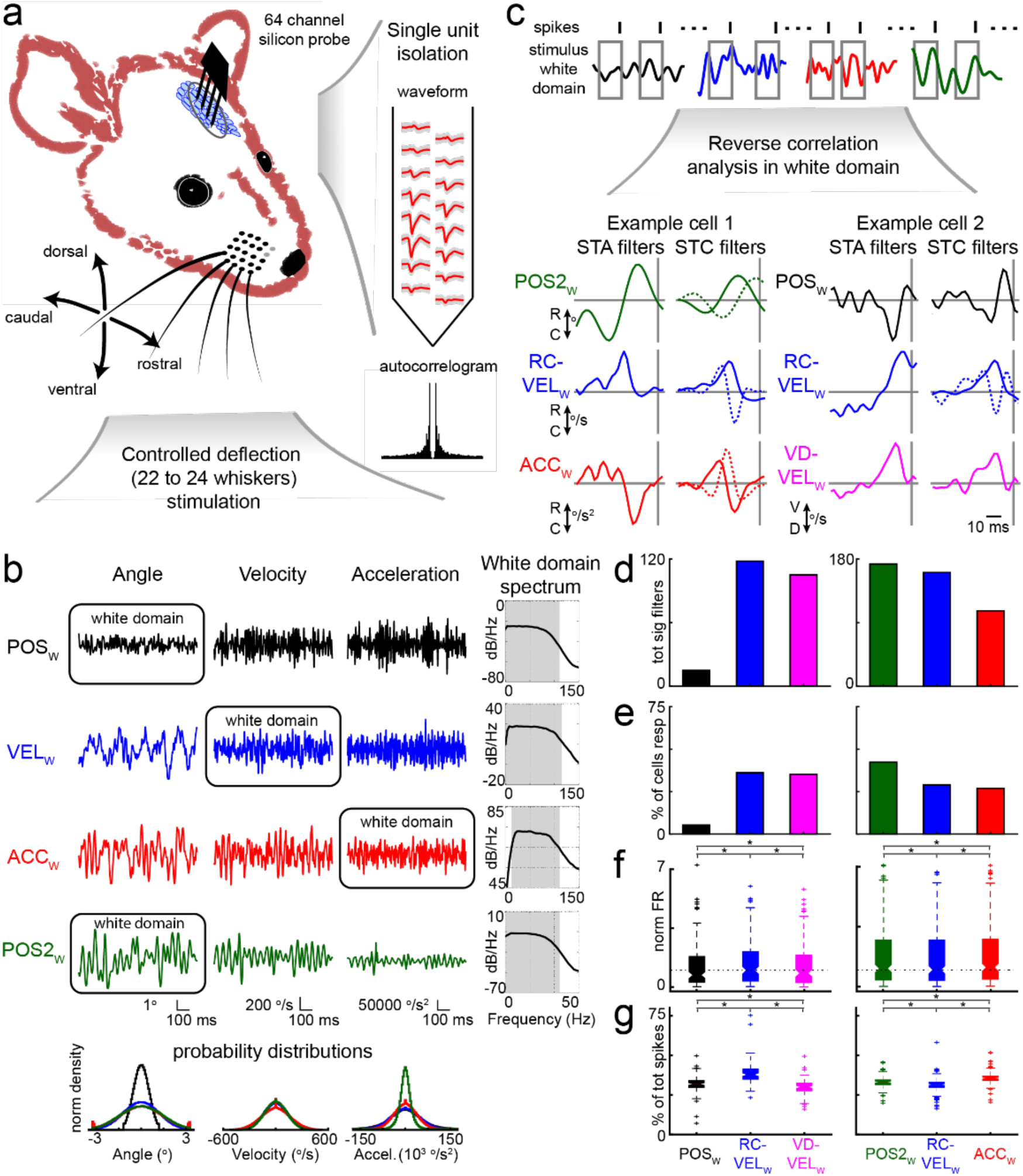
Neuronal responses under stimulations optimized for studying distinct kinematic features (position, velocity, acceleration). **(A)** Schematic of the experimental approach. A two dimensional piezo-electric controlled stimulation is applied to 22 to 24 whiskers (only four displayed, bottom). Simultaneous recording of hundreds of neurons was made with 64-channel silicon probes across all layers of wS1 (right). **(B)** Stimulus portions of POSW, VELW, ACCW and POS2W and kinematic properties (angle, velocity, acceleration). Their white domain spectral content (gray shaded, right), and their probability distributions across stimuli (bottom). **(C)** The reverse correlation method collects short windows before the spikes for each stimulus white domain to compute a Spike Triggered Average/Correlation on them (STA/STC, top). Example STA and STC filters obtained from two cells. Only STC filters in velocity are single sided (bottom). **(D- E).** Two sets of experiments were performed with three stimuli each, repeating RC-VELW for proper comparison (batch 1, left column, 151 cells, batch 2, right column,254 cells). **(D)** Number of filters obtained for each stimulus. **(E)** Percentage of significantly responding cells. **(F)** Normalized firing rates (Rank Sums test, p<0.001). **(G)** Percentage of spikes across stimulus (Rank Sums test, p<0.001).

An effective stimulus should strategically sample the relevant dimensions of the stimulus space to uncover neuronal coding, without requiring exhaustive exploration. At the same time, it needs to temporally separate the relevant features encoded by the neurons to allow their selective contribution to be isolated during the analysis. According to the stick/slip hypothesis (16), a high velocity and/or acceleration stimulus will elicit the higher number of cortical spikes. However, according to the sweep/stick hypothesis (15), only high velocity stimuli, and not high acceleration stimuli, are sufficient to reveal feature selectivity in wS1 neurons, going beyond simply eliciting spikes. This limitation likely contributed to a reduced firing rate, as observed previously (15). Based on this, we designed stimuli with matched velocity distributions (Fig. 1B, middle column) and distinct profiles for position (POS_W_, black), velocity (VEL_W_, blue) and acceleration (ACC_W_, red), each sampled from white noise Gaussian distributions (Fig. 1B, white domains). For POS_W_, it is not possible to generate the same angular distribution as for VEL_W_ and ACC_W_ due to the nature of white noise in position, which inherently results in smaller deflection amplitudes. This limitation likely contributed to a reduced firing rate, as observed previously (15). To address this, we introduced a second position-based stimulus (POS2_W_, green) with broader angular excursions. This came at the cost of reduced acceleration range and a flat spectrum limited to 35 Hz in the white domain (Fig. 1B, right column). Additionally, we applied a ventro-dorsal velocity stimulus (VD-VEL_W_) to explore potential 2-dimensional stimulus preferences.

To determine the relevant stimulus characteristics that neurons respond to, we used reverse correlation methods such as spike-triggered averaging (STA) and spike-triggered covariance (STC). These methods allow to extract the features (filters) that are encoded by a given neuron by analyzing small time-windows of spike-eliciting stimulus events (Fig. 1C). Two batches of experiments were done, one with POS_W_, RC-VEL_W_ and VD-VEL_W_, stimulations and another with POS2_W_, ACC_W_ and RC-VEL_W_, stimulations repeating RC-VEL_W_ stimulation in both batches for better comparison (see Methods). We measured individual neuronal responses in 405 cells on 7 animals. In the two example cells shown in Fig. 1C, only RC-VEL_W_ and VD-VEL_W_ have a monophasic STC filter, characterized by a single sided deflection in time, indicating a preference for unidirectional stimulus events. In contrast, other stimuli display biphasic or oscillating filters, which include both positive and negative lobes, suggesting sensitivity to stimulus changes or bidirectional movement patterns. Therefore, we can parsimoniously assume that these neurons are encoding for a unidirectional deflection of the whiskers in response to a velocity stimulus event. Our following analyses are based on the STC results and not the STA, since most of the neurons have an STC and do not present an STA (see Methods, see also (17,18) for similar observations).

To compare the efficacy of each stimulus in activating the neurons, we looked at the significant filters for all the recorded neurons. In the first batch of experiments (151 cells, left column in Fig. 1D-G), the position stimulus (POS_W_, black) showed a substantially reduced number of filters compared to the rostro-caudal velocity stimulus (RC-VEL_W_) (Fig. 1D). Also, stimulation in position (POS_W_) rendered a lower number of responding neurons (Fig. 1E), less firing rate (Fig. 1F) and less relative number of spikes across stimuli (Fig. 1G). The ventro-dorsal velocity stimuli (VD-VEL_W_, pink) also elicits less spikes, firing rate, and renders less significant filters than RC-VEL_W_. However, the percentage of cells responding with feature selectivity (displaying filters) is almost as much as with RC-VEL_W_.

For the second batch of experiments (254 cells, right column in Fig. 1D-G), the POS2_W_ stimulus (green), contrary to POS_W_, allowed us to find more significant filters than the RC-VEL_W_. This can be partly explained by the stimulus design. Since the white noise spectrum is flat up to 35Hz, the stimulus correlation across time is higher, and the significance threshold is easier to cross to find significant filters (a smaller number of spikes needed for an equivalent recording time, see Methods). In this batch, the POS2_W_, and the ACC_W_ (red) stimuli have more firing rate and spikes compared to the RC-VEL_W_, but ACC_W_ produces less significant filters, and less responding cells. The increase in spike count is probably caused by the slight increase of high position values explored by these two stimuli (see the edge of the angle probability distribution in Fig. 1B, bottom, see also the Methods). However, at least for the ACC_W_, these spikes are not as relevant as RC-VEL_W_ induced spikes for the neurons because they produce less significant filters.

### A low dimensional representation of neuronal stimulus preference

We extracted the most relevant features at the population level using a principal component analysis (see Methods). In other words, for each stimulus white domain, we studied if the filters encountered for all the neurons could be represented by a few representative filters. We found that for each stimulus, almost 90% of all the significant filters could be explained by two principal components (PCs) (Fig. 2A, left). We can then create a feature space using these two filters as two orthogonal axes. In these feature spaces obtained for each stimulus white domain, each neuron significant filter can be plotted as a linear combination of the two PCs (Fig. 2A, right). The closer to the unit circle the cell’s filter is, the better represented by the PCs that create the subspace. For all the stimulus subspaces there is a strong tuning to the unit circle. Remarkably, the only subspace showcasing a specific organization is the RC-VEL_W_. The neuronal population of significant filters are tuned to specific regions of this subspace, which correspond to specific whisker deflections. We followed to study this organization in detail because it reflects the encoding of the whisker system at the level of the barrel cortex. A similar organization in the VD-VEL_W_ stimulus is present but it’s less well defined (see Supplementary Fig. S2).

**Figure 2.**
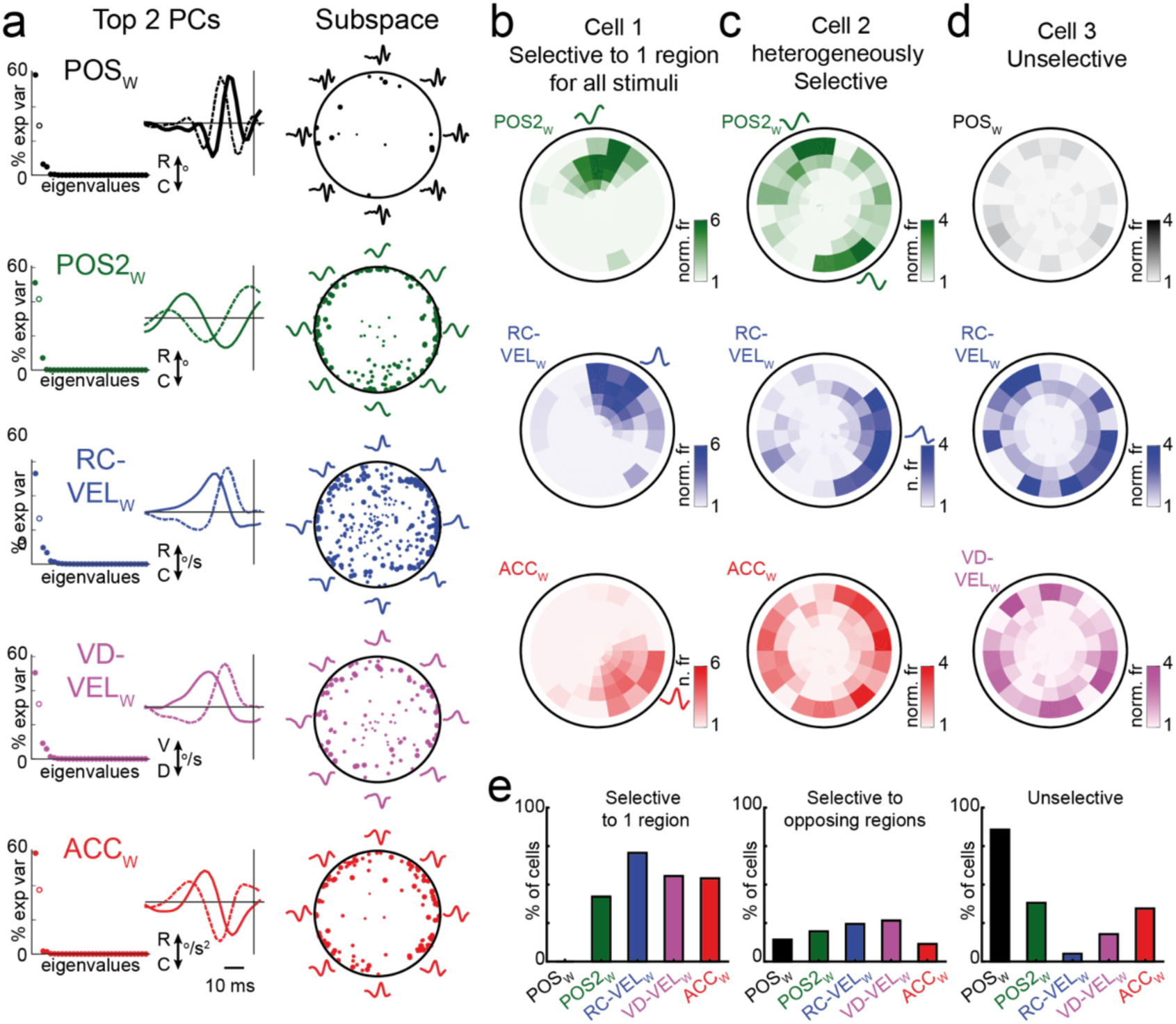
Filter spaces obtained for distinct kinematic features. **(A)** Response subspaces obtained by principal component analysis made on the filters obtained for the five different types of stimulation presented. Left: Variance explained. Center: First (bold) and Second (dashed) PCA components. Right: Projection of all significant filters from all recorded cells onto the two-dimensional subspace defined by the population PCA. Large dots for first and second, and small for third and fourth significant filters of each cell. All subspaces capture almost 90% of the variance of the filters of each stimulus. A higher number of filters and a segregation in filter space is most apparent for RC-VEL_W_ stimulation. **(B-D).** Example of tuned cells, showing each cell’s firing-rate tuning within the population feature space (not its filters). Cells show selectivity through an increased firing rate to specific regions near the unit circle of the subspace:: **(B)** Selective to one region **(C)** Selective to two opposing regions (top), one region (middle) or unselective (bottom). **(D)** Unselective. **(E)** Percentage of cells selective to one region (left), opposing regions (middle) and unselective (right).

We then determined how each cell is tuned in the feature spaces. For this, we plotted their firing rate in different angular and radial portions (Fig. 2B-D). In this representation, a cell can be selective to one filter shape (Fig. 2B), two opposite filter shapes (Fig. 2C) or it can be unselective (Fig. 2D), meaning that it responds equally to any shape near the unit circle. To test for significance, we did two shuffle statistical tests for unidirectional and bidirectional tuning (see Methods). By analyzing the cell population, each stimulus can be characterized by its selectiveness (Fig. 2E). The RC-VEL_W_ stimulus is the most selective to one region of the feature space, the VD-VEL_W_ stimulus is the most selective to opposing regions, and finally the RC-VEL_W_ stimulus is the most selective, considering both possibilities. These results confirm our hypothesis that the RC-VEL_W_ stimulus should be the most effective for identifying more precisely the information coded in wS1. In the following results, we will study in-depth the neuronal responses to this specific stimulus.

### A sweep-stick coding across the different layers of wS1

In the RC-VEL_W_ feature space (Fig. 2A, right), we can define a sweep/stick axis (Fig. 3A), as was done previously in the rat S1 (15). The sweep and stick feature shapes are similar in both species (see Supplementary Fig. S3). When we plot the density of neurons significantly tuned to specific angles in this space (Fig. 3B), the distribution shows two peaks closely corresponding to the sweep and stick axes. To classify neurons into each coding class, we used a method defined previously in rats on neurons with significant responses using a sparse noise stimulation across the 24 whiskers (see Methods). The “stick” cells are fast single-whisker encoding cells, while the “sweep” cells are slow multi-whisker encoding cells. We obtained 21 stick-, 52 sweep- and 13 mixed-responding cells (Fig. 3C) from the 405 recorded cells (the low yield is because cells need to show significant responses to both for RC-VEL_W_ and sparse noise stimuli, the latter being less effective than the white noise stimulus). We plotted the stick and sweep density on the stick/sweep feature space (Fig. 3D) and found that the stick and sweep-responding neurons accumulate around the two axes defined above. Therefore, the classification of neurons based on the stick and sweep feature response selectivity matches the one defined previously in rats (15).

**Figure 3.**
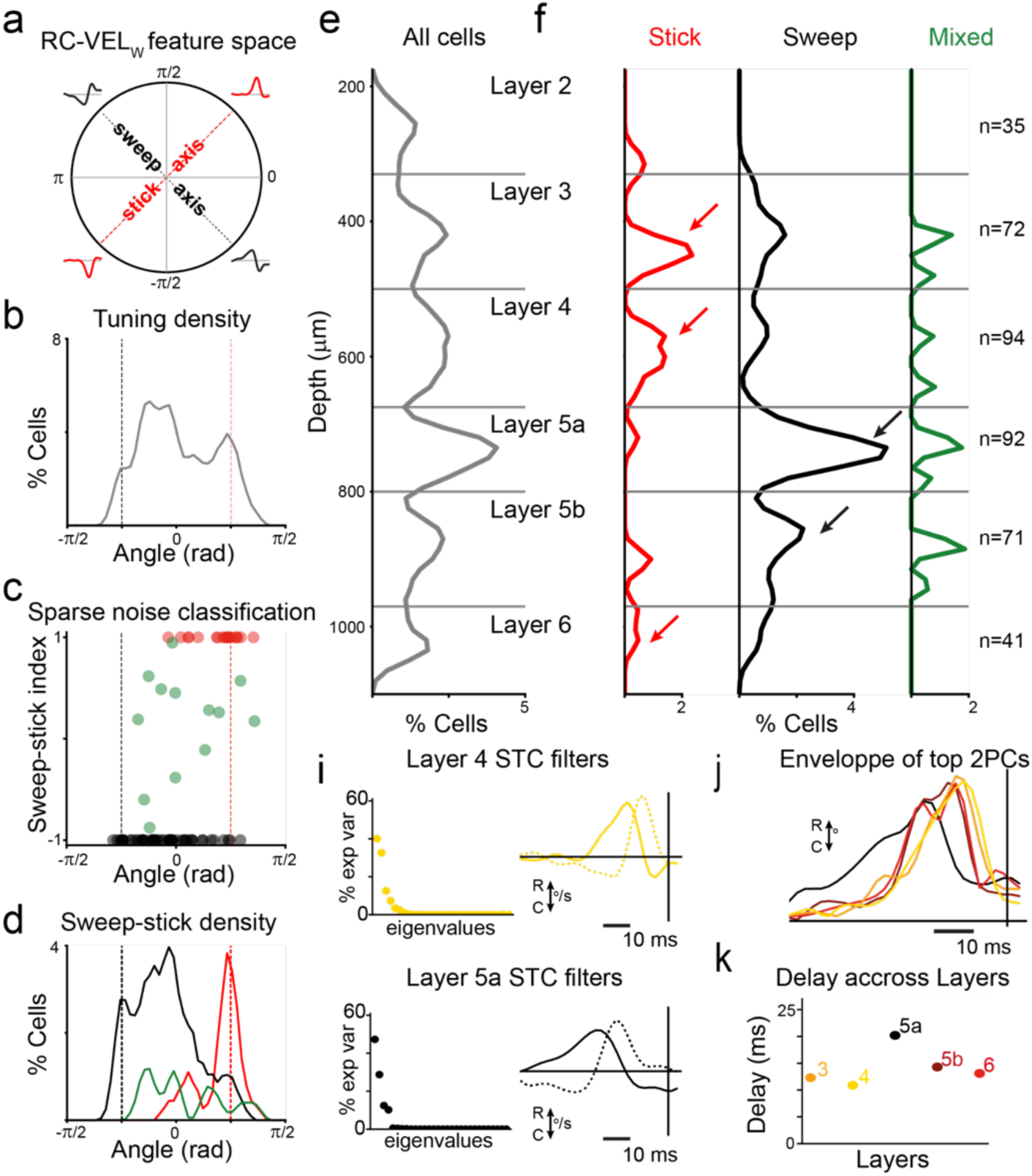
The sweep-stick coding in the barrel cortex is layer dependent. **(A)** Sweep-stick feature space. **(B)** The density of cells tuned to a specific angle in the Sweep-stick feature space. **(C)** Classification of sweep (n=52, black), stick (n=21, red) and mixed (n=13, green) -responding cells using a Sparse Noise stimulation sweep-stick index. **(D)** Distribution of sweep (black) stick (red) and mixed (green) -responding cells across the Sweep-stick feature space. **(E)** Distribution of all the cells recorded across the layers in the barrel cortex. **(F)** Distribution of each cell class in the layers of the barrel cortex. Cell counts of each class in each layer (right). **(I)** Principal component analysis made on the filters of layer 4 and 5a. Left: variance explained, Right: first (bold) and second (dashed) PCAs component. **(J)** Energy envelope made by the top two PCs for each layer. **(K)** Peak delay (latency) of the envelope for each layer (Delay per layer: L3: 12.3 ms, L4: 10.9 ms, L5a: 20.2 ms, L5b: 14.3 ms, L6: 13.1 ms).

Unlike previous works using STC analysis, where the number of spikes needed to extract significant filters is high (18), we succeeded here at the supra-granular cortical layers of wS1 where spiking activity is relatively low to obtain many responding neurons (15). This was due to the use of the optimized stimulus spaces. Thus, we could explore if “stick-” and “sweep-” functional classes are segregated in different layers of the barrel cortex. “Stick” cells were found mostly in layers 3, 4 and 6, whereas “sweep” cells were registered mostly in layers 5a and 5b. Mixed cells are homogeneously distributed from layer 3 to layer 5b (number of measured cells per layer: L2: 35, L3: 72, L4: 94, L5a: 92, L5b: 71, L6: 41, Fig. 3G). We found a specific organization of “stick-” and “sweep-” cells across layers, with a repartition for the granular and infragranular layers equivalent to that previously shown in the rat (Fig. 3F) (15).

We then analyzed the resulting filters from the STC analysis across layers. For this, we computed a principal component analysis (PCA) on the filters encountered in neurons of layer 4 and 5a, which have a peak in the stick and sweep-responding cells density, and we extracted the first two principal components (PCs) (Fig. 3I). The PCs shapes in layer 5a are more delayed and extended compared to the ones in layer 4, suggesting that temporal latency might play a role in distinguishing sweep and stick feature selectivity. Specifically, longer delays may reflect the need to integrate information over longer time windows, which could be necessary for encoding sweep movements that evolve more gradually over time, as opposed to the more abrupt stick events. To quantify this, we computed the PCs subspace envelope of each layer, which represents the subspace energy contribution of the filters before the spike time (Fig. 3J, see Methods). We obtained the time of the peak energy before the spike for each envelope to compare the delays between layers (L3: 12.3 ms, L4: 10.9 ms, L5a: 20.2 ms, L5b: 14.3 ms, L6: 13.1 ms, Fig. 3K). According to this measure, the whisker deflection information arrives first in layer 4 and lastly to layer 5a. The longer delay and broader envelope in layer 5a suggest that neurons in this layer integrate inputs over a wider temporal range, likely enabling them to extract features from longer, more complex whisker deflection patterns. As mentioned above, layer 5a receives information from POm in the thalamus via the paralemniscal pathway. This pathway principally innervates layer 1 and 5 of wS1, and the lemniscal pathway connects the VPM in thalamus mainly to layer 4 and 6 of wS1. The distribution of “stick” and “sweep” responses in wS1 is in line with these pathways’ organization (higher stick-responding cell density in layer 4 and 6, and higher sweep-responding cell density in layer 5a). The next question we asked is whether this cortical sweep/stick encoding is inherited from these nuclei of the thalamus?

### wS1 “stick” coding is inherited from the thalamus in a layer specific manner

To explore if the sweep/stick cortical coding scheme is inherited from the thalamus, we recorded neurons in the thalamic nuclei VPM and POm, using silicon probes during white noise simulations of rostro-caudal velocity-optimized deflections of whiskers (VEL_W_, Fig. 4A). We recorded 105 cells from POm and 43 cells from VPM (the histological position of the recorded cells can be seen in Fig. 4B and Supplementary Fig.4), to which we applied our reverse correlation analysis. As for wS1 cells, we extracted the significant filters from the ensemble of spike-eliciting stimuli for each cell using STA and STC analyses (see Fig. 4C for two example cells). Then we applied a principal component analysis on the significant filters separately for each structure (Fig. 4D). Here again, as in wS1, more than 80% of the variance could be explained by the first two PCs. However, compared to the cortex, the first PC eigenvalue has a higher relative weight than the second one. We also corroborated that RC-VELw is the most adequate stimulus to study feature selectivity in these cells (Supplementary Fig. S5). The shapes of the first two PCs are similar for VPM and POm and also resembled the first two PC shapes of wS1. This structured distribution of filter types across thalamic neurons suggests that the thalamus itself may implement a form of sweep/stick encoding, which is then relayed to the cortex in a nucleus, and layer-specific manner.

**Figure 4.**
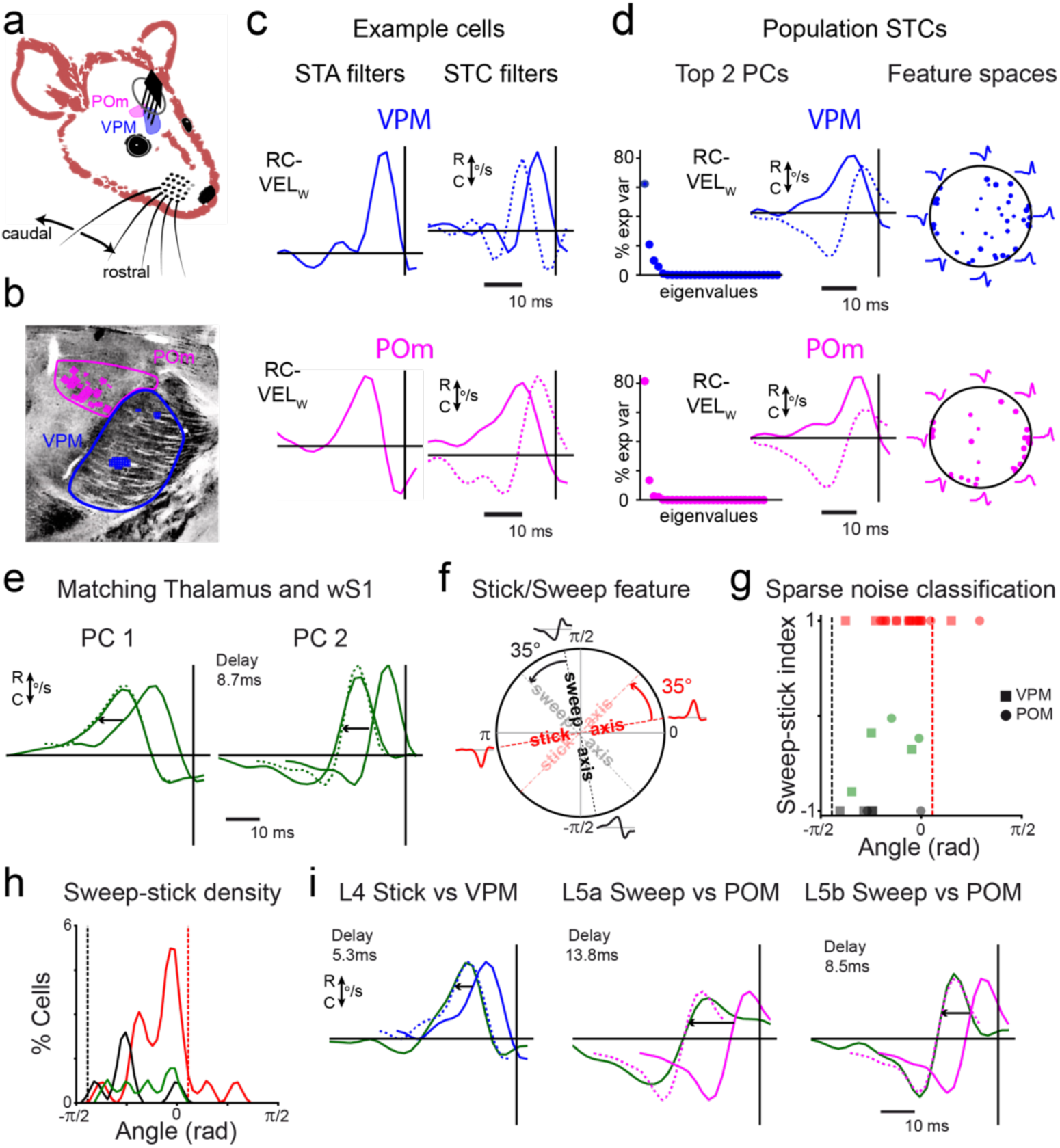
Thalamic sweep-stick features encoding is inherited by the barrel cortex. **(A)** A 64 channels silicon probe is placed in the thalamus of a mouse during a caudo-rostral whisker stimulation **(B)** Histological reconstruction of the position of each recording site in VPM and in POM. **(C)** Two cell examples in VPM and in POm with their STA and significant STCs. **(D)** Response subspaces obtained by principal component analysis made on the filters of POm (n=105) and VPM (n=43). Left: Variance explained. Center: First (bold) and second (dashed) PCAs components. Right: Projection of all significant filters from all recorded cells onto the two-dimensional subspace defined by the population PCA. **(E)** Matching between the first two PCAs components of thalamus (148 cells) and wS1 (405 cells) is done by rotating and delaying the subspace PCAs of thalamus. The best match has a 35° rotation and an 8.7ms delay. **(F)** Stick-Sweep feature space in the thalamus, obtained by a 35° rotation of wS1 feature space. **(G)** Classification of sweep (n=6, black), stick (n=21, red) and mixed (n=5, green) -responding cells across the Sweep-stick feature space for VPM (square) and POM (circle) using a sparse noise stimulation. **(H)** Distribution of sweep (black), stick (red) and mixed (green) -responding cells across the Sweep-stick feature space. **(I)** Matching by rotating and delaying one PCA feature of VPM (n=43) or POm (n=105) to the stick and sweep features of specific layers in wS1 (L4: 94 cells, L5a: 92 cells, L5b: 71 cells).

To assess whether the cortical sweep/stick feature space could be inherited directly from thalamic representations, we first combined the data from VPM and POm into a single thalamic population. This strategy was motivated by the similarity of their PC shapes and the lack of clear separation in their feature encoding (as shown above), suggesting they could share a common encoding subspace. We then performed a new PCA across all 148 thalamic neurons and compared the resulting top two PCs to those obtained from the cortical neurons. To do so, we rotated the thalamic PC space and applied a temporal shift to find the best match to the cortical PC subspace. Functionally, this rotation corresponds to a change in how feature selectivity is aligned across populations: a rotation of the subspace reflects a re-weighting of sensitivity to different stimulus dynamics (e.g., timing or phase of motion), while a delay captures transmission or integration time between structures. The best rotation and delay values were determined by maximizing the total scalar product between the delayed and rotated thalamic PCs and the cortical PCs (see Methods). The best match corresponds to a 35° rotation angle and 8.7 ms delay (Fig. 4E). Another way to illustrate this match is in the sweep-stick axis defined for wS1 as shown in Fig. 4F: the thalamic feature space needs to rotate 35° to match the cortical subspace. Conversely, this result allows us to define the stick and sweep axis in the thalamus by decreasing them by 35° (dashed lines in Fig. 4G). Using the same sparse noise classification as in the cortex, we found stick and sweep-responding neurons in POm and VPM (Fig. 4G). If we plot the sweep/stick density in the feature space, we observe that the two populations are following the two newly defined thalamic sweep and stick axes (Fig. 4H), showing that our sweep/stick classification is also present in the thalamus, with the sweep-responding cell density tuned closer to the stick axis than in wS1. The cells in VPM and POm are mainly of the “stick” functional class, debunking the hypothesis of a segregation between stick and sweep coding in these two nuclei.

To better understand the respective contributions of the lemniscal (VPM) and paralemniscal (POm) pathways to cortical coding,, we compared the thalamic feature subspaces to those of cortical layers in wS1 (Fig. 4I). Specifically, we quantified the best match between the first two principal components (PCs) of VPM and POm and those of each wS1 layer (L3 to L6). We found that the VPM subspace aligned best with the stick-coding subspace of layer 4, with the shortest delay of 5.3 ms. This strongly suggests a direct and fast transmission of stick feature coding from VPM to layer 4. On the other hand, when matching POm feature subspace with the sweep-coding subspace of layer 5a, this was poorer and associated with a substantially longer delay. This indicates that while POm may contribute to the sweep feature representation in layer 5a, it cannot fully account for the emergence of this coding pattern. As an additional comparison, we found that POm feature subspace also aligns reasonably well with layer 5b, with a second-best match and only a 8.5 ms delay, again shorter than the match with 5a. These results suggest that, unlike stick coding which appears to be directly inherited from VPM to layer 4, sweep coding may emerge through intracortical processing, potentially integrating POm inputs with stick-like signals from layer 4. This supports a model where layer 5a acts as a site of feature recombination or transformation, rather than a simple relay of thalamic input.

### Caudal/rostral tuning of stick/sweeps-responding cells allow wS1 differential feature encoding

To better understand the relation between sweep and stick coding and have a better description of the emergence of sweep coding in wS1, we did a further characterization of the neuron coding features. We classified neurons into fast and regular spiking cells using their extracellular spike-waveform. This is a proxy for identifying excitatory and inhibitory cells (Fig. 5A, see Methods) (19). We found that 76% of the cells (328 cells) are regular spiking and 19% are fast spiking cells (81 cells), and 5% were unclassified units (“undefined”, 20 cells). In the thalamus, the cells were regular spiking (Supplementary Fig. S6). Then we studied the distribution of fast and regular-spiking neurons, having sweep and stick coding properties across cortical layers (Fig. 5B). Sweep-responding cells are predominantly regular spiking cells, and stick-responding cells, especially in layer 3 and 4, are mostly fast spiking cells. In layer 4 and 5a, we find mostly that fast spiking neurons belonged to the stick functional class and regular spiking neurons to the sweep functional class respectively. We asked then, do these groups have the same velocity directional preference in the caudo-rostral axis? By examining directional tuning, we aimed to determine whether the stick and sweep-responding neuronal populations not only differ in temporal processing and cellular identity but also encode distinct spatial features of whisker movement, such as direction along the caudo-rostral axis. This could further clarify the functional divergence between these populations and their respective roles in sensory encoding within wS1.

**Figure 5.**
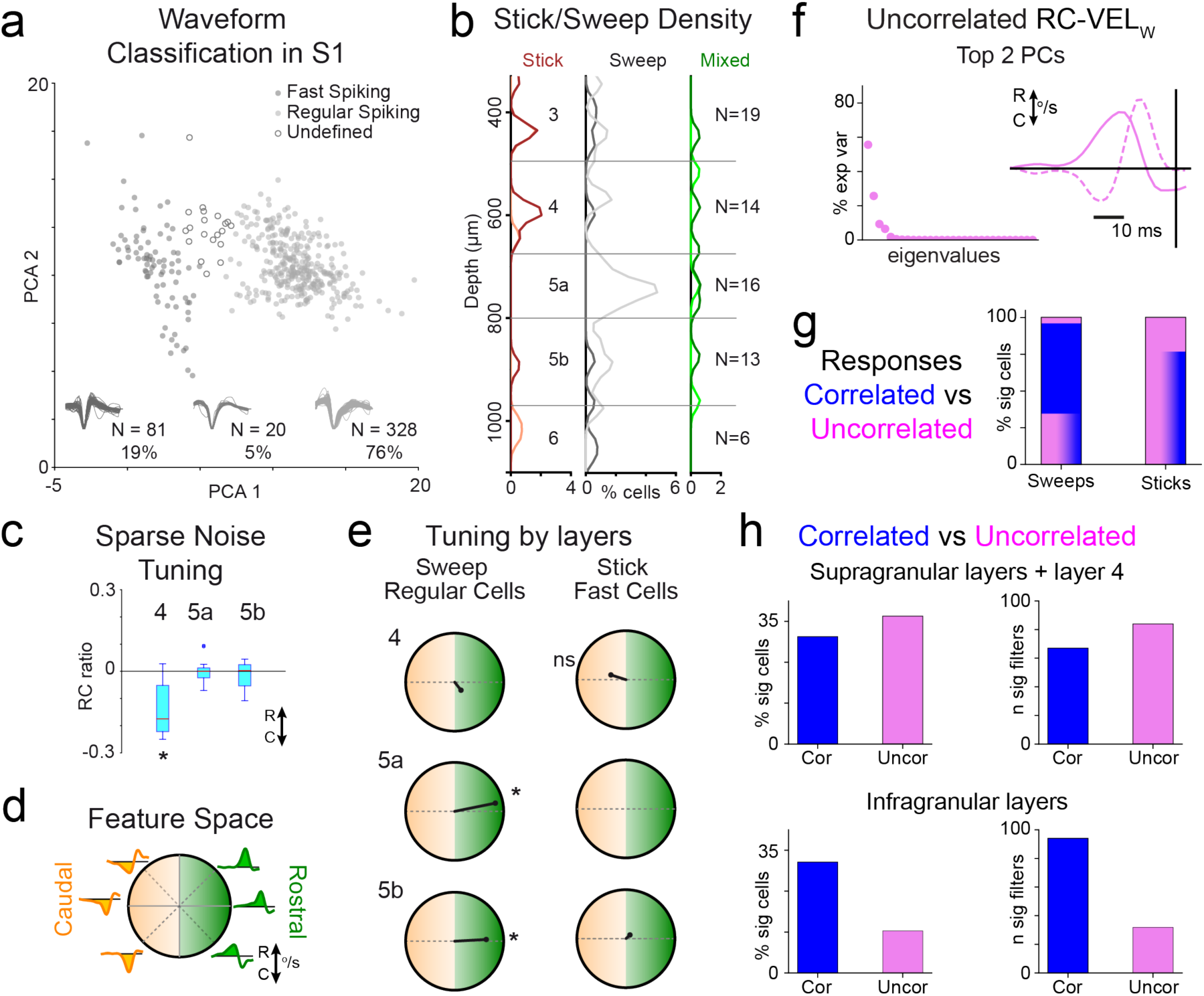
Caudal / rostral and correlated / uncorrelated tuning across wS1 layers of fast-spiking stick-responding neurons and regular-spiking sweep-responding neurons. **(A)** Spike-waveform classification in wS1 of fast (dark grey) and regular (light grey) spiking cells. **(B)** Density of each cell type across layers. Fast (dark red) and regular stick-responding neurons (light red); Fast (dark grey) and regular sweep-responding neurons (light grey); Fast (dark green) and regular (light green) neurons with mixed responses. **(C)** Sparse noise classification of rostro-caudal preference for layer 4 fast stick-responding cells (n=5) and regular sweep-responding cells of layer 5a (n=13) and 5b (n=8). Fast stick-responding cells in layer 4 are significantly tuned to caudal movements (p<0.05). **(D)** Caudo-rostral tuning in the Sweep/Stick feature space. **(E)** Tuning by layers for regular sweep and fast stick-responding cells. Lines represent the vector sum of all tuned cells in the corresponding layer. Layer 5a and 5b are significantly tuned rostrally (directional shuffling test, p<0.02). **(F)** Response subspaces obtained by principal component analysis made on the filters of wS1 (n=116). Left: Variance explained. Right: First (bold) and second (dashed) PCA components. **(G)** Percentage of significantly sweep- and stick- responding cells to correlated (blue), uncorrelated (pink) or both (blue-green blend) RC-VEL_W_ stimuli. **(H)** Percentage of significantly responding cells (left) and number of significant filters (right) to correlated (blue) and uncorrelated (pink) RC-VEL_W_ stimuli, for supragranular plus layer 4 (top) and infragranular layers (bottom).

To test this, we used a sparse noise stimulus to calculate rostro-caudal tuning ratios (RC-ratio) by counting the spikes corresponding to forward or caudal single whisker velocity movements (see Methods) for the cells in layer 4, 5a and 5b (Fig. 5C). The RC-ratio showed significantly caudally tuned fast spiking stick-responding cells in layer 4 (Wilcoxon test, p<0.05). On the contrary, this stimulus did not result in any tuning preference for the regular spiking sweep-responding cells of layer 5a and 5b. Since our sparse noise stimulus deflects only one whisker at a time, it may not sufficiently engage sweep-responding cells, which are potentially more responsive to multi-whisker or spatiotemporally complex stimuli. Therefore, in our velocity feature space obtained during RC-VEL_W_, we defined a rostral (green) and a caudal (orange) deflection region (see Methods, Fig. 5D). We computed the tuning for each group of cells by calculating the population vector as a sum of their direction preference in this space. We found that the regular spiking sweep-responding cells in layer 5a and 5b are significantly tuned rostrally (directional shuffling test, p<0.02, Fig. 5E). As expected, the fast-spiking stick-responding cells show a tendency toward caudal tuning in this feature space, although this was not statistically significant, consistent with the idea that they are best engaged by single-whisker deflections as in the sparse noise condition.

### Sweep-responding cells respond to correlated whisker movements in lower layers, while stick-responding cells respond to uncorrelated ones in upper layers

Having characterized directionality preference across layers by sweep/stick functional classes, we next asked whether uncorrelated dense whisker movements would reveal a finer feature code. This had previously been unattainable: a 24-dimensional increase in the stimulus requires a corresponding 24-fold increase in spikes to obtain detectable responses. However, the low-dimensional subspaces we recovered from correlated stimuli suggested that an uncorrelated RC-VEL_W_ stimulus could resolve finer feature subspaces even at the low spiking levels of the upper layers, demonstrating again the value of an optimized stimulus space for probing responses to complex stimuli. Analyzing the same cells as in the correlated condition, we obtained a similar two-dimensional PC subspace explaining almost 90% of the significant filters (Fig. 5f). Two example cells (stick- and sweep-responding) with their significant filters across the whisker pad are shown in Supplementary Fig. S7a-b. As expected, sweep-responding cells were mostly tuned to correlated stimuli (Fig. 5g, left: 61% correlated only, 32% both, 7% uncorrelated only), while stick-responding cells were mostly tuned to uncorrelated stimuli (Fig. 5g, right: 100% tuned to uncorrelated, of which 80% were also tuned to correlated). By cortical depth, upper layers (supragranular plus layer 4) yielded more significant filters and more cells responding to uncorrelated stimuli (Fig. 5h, top), whereas infragranular layers were dominated by filters and cells responding to correlated stimuli (Fig. 5h, bottom).

In some stick-responding cells we recovered significant filters in neighboring whiskers. Repeating the subspace tuning analysis of Fig. 2 for each whisker independently (Supplementary Fig. S7a-b, right for two cell examples) revealed a further difference between cell classes: in stick-responding cells the principal (PW) and the neighbour secondary (SW) whiskers tend to be tuned to opposing regions of the VEL_W_ subspace, while sweep-responding cells show a flatter angular difference. Across the population, the distribution was enriched near a 180° PW-SW tuning difference (Supplementary Fig. S7c). Whereas separating the populations, this enrichment was present only for stick-responding and mixed-responding cells. This antialignment in stick-responding cells provides further evidence for a specialization of detailed single-whisker features in the upper layers, while features encoded by sweep-responding cells, tuned to more correlated multi-whisker movements, are encoded in the lower layers.

### A cortical sweep-responding neuron model integrates thalamic stick-responding cell information

The results shown until now provide a description of how different selectivity features such as stick/sweep, caudal or rostral, correlated or uncorrelated preference are distributed in the different regular and fast spiking cell types across the layers of wS1 and within thalamic nuclei VPM and POm. Here we propose a mechanism by which all these properties may come together in the somatosensory pathway to allow an animal to recognize objects and textures in its environment when it is actively whisking (Fig. 6A).

**Figure 6.**
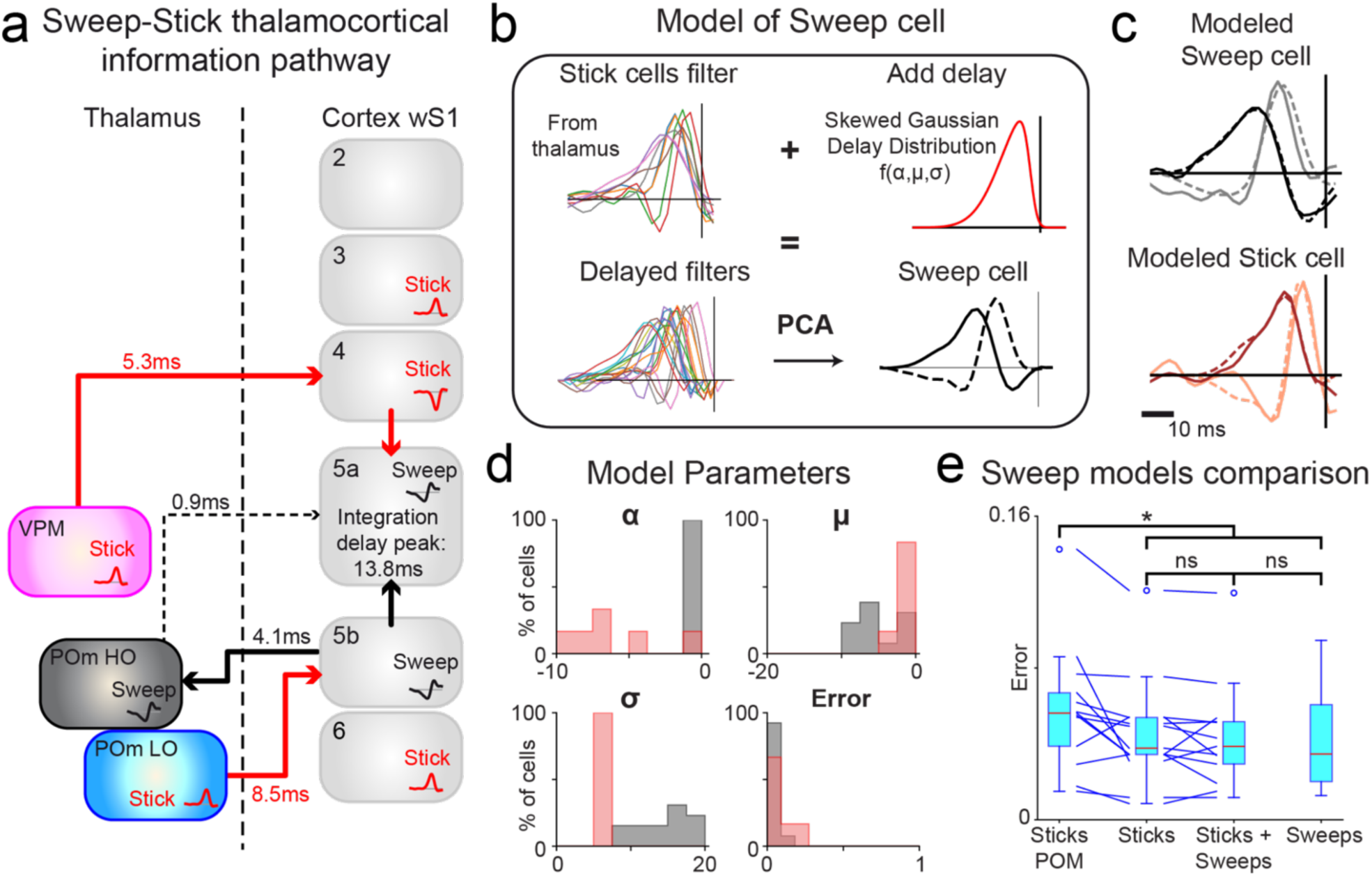
A model with time integration of thalamic stick filters gives rise to sweep- and stick-responding neurons in the cortex. **(A)** Model of whisker information propagation from the thalamus to the cortex wS1. Rapid stick-responding cells caudal inhibition may produce brief local circuit silencing, allowing signal sharpening in layer 4 after single whisker contact detection. Delayed rostral integration of multi whisker information may help collect layer 4 (VPM relayed), and POm stick information, giving rise to the more complex sweep encoding in deep cortical layers, thus allowing for texture feature coding. Higher order (HO) POm presents sweep information with a latency similar to layer 5a, inherited from layer 5b (see Fig. 7). Given the motor and secondary somatosensory inputs to HO POm, and its strong innervation of layer 5a, it may modulate the sweep code arising in layer 5a. **(B)** Model of Sweep-responding cells. We add delays and augment multiple stick filters from VPM and POm to create a filter ensemble. We extract the two model sweep filters by doing PCA. Each sweep-responding cell from layer 5a was modeled by fitting the three parameters of a skewed Gaussian delay distribution. **(C)** Top: Model of one sweep-responding cell of layer 5a created with stick-responding neurons from POm and VPM. Bottom: Model of one stick-responding cell of layer 4 created with stick inputs from VPM. (1st filters: black, 2nd filters: gray, model: dashed lines, experimental: solid lines). **(D)** Histograms of the fitted parameters for the two models: stick-responding cells of layer 4 (n=6) modeled by stick- responding cells of VPM (n=10) and sweep-responding cells of layer 5a (n=13) modeled by stick- responding cells of VPM and POm (n=17). **(E)** Distributions of errors for four models of sweep- responding cells from layer 5a created with i) Stick-responding neurons from POm (n=7), ii) Stick- responding neurons from POm and VPM (n=17), iii) Stick- and Sweep-responding neurons from POm and VPM (n=22), iv). Sweep-responding neurons from POm and VPM (n=5). Paired-t test, model 1 vs model 2 (p<0.05), model 1 vs model 3 (p<0.01), model 1 vs model 4 (p<0.02).

When a mouse whisks, it deflects its whiskers back and forth in the rostro-caudal direction at frequencies between 15 to 30 Hz (20). When the animal encounters an object, many whiskers make contact with it during the protraction whisking phase. These contact events happen at different times, creating multiple fast caudal stick-type of movements. This information arrives first in layer 4 and layer 5b, from VPM and POm respectively. The fact that many stick-responding cells in layer 4 may be inhibitory hints to the fact that they may act as a reset of the ongoing local responses in the regular spiking cells from self-whisking movement, they may act as a way to sharpen in time the local circuit responses (21,22). Further fine spatial tuning in stick-responding cells is provided by the anticorrelated velocity tuning between the PW and SW. When a whisker encounters a small object or edge, the neighboring whisker deflects in the opposing direction, allowing better spatial localization of the first stick movement.

Once the whiskers have come in contact with the surface of the object, the animal will continue for a brief period of time to protract them. If the surface is not smooth, the whiskers may encounter an irregularity where they may get stuck, to then be released together almost at the same time. This release will consist on a multiwhisker high velocity rostral deflection event, which will happen at slightly different moments, i.e. a wider time window than the single whisker stick event. If we consider that this happens only during a quarter of the whisking cycle, from the midpoint angle range to the maximal rostral position, the maximal time window of this event will be of ∼16 ms. The whiskers will have made a sweep-like movement release, in a short but not instantaneous time window, from a texture present on the surface of the object. We propose that this type of information will be integrated in layer 5a neurons, from the information coming from layer 4 or layer 5b. But is it possible to create sweep-like response features from the information gathered from stick-responding cells activity?

We hypothesize that sweep-like responses are generated in the cortex by integrating in time multi-whisker information from multiple stick-responding cells. To test this, we built a model for the emergence of sweep-responding cells by integrating stick-responding cell information in a time window using our measured stick-type filters (Fig. 6B). We selected the first filter of all the stick-responding cells from thalamus (VPM and POm), and we added a delay to them to mimic how this information would arrive at layer 5a. We also did data augmentation to obtain between 750 to 1700 filters (see Methods), which corresponds roughly to the number of spikes recorded for the sweep-responding cells. To allow for physiological delays we selected them from a skewed Gaussian distribution (see Methods). These filters are our modeled spike triggered ensemble, to which we can apply principal component analysis to extract the filters of a model sweep-responding cell (Fig. 6B, bottom). We fitted the parameters of the skewed Gaussian so that the filter curves minimize the difference (error measure, see Methods) between our measured sweep-responding cell filter and the modeled one. We modeled this way all the sweep-responding cells filters from layer 5a (n=13) with all the stick-responding cells from POm and VPM (n=17, example is shown in Fig. 6C, top). This model is versatile, and allowed us to also model the stick-responding cells (n=6) from layer 4 using only the stick-responding cells from VPM (n=10) (an example is shown in Fig. 6C, bottom). To compare the difference between the sweep and stick-responding modeled cells, we looked at the distribution of the fitted parameters (Fig. 6D). The mean delay of the stick-responding cells is smaller than for sweep-responding cells. But more remarkable is the difference in the standard deviation of delays, where sweep-responding cells show a much longer distribution. In this way we confirm that sweep-responding cells filters of layer 5a can be obtained by gathering stick-responding cells information in a relatively wide time window. But, is the integration of information from both VPM and POm stick-responding units really necessary to obtain sweep responses?

We therefore compared the sweep-responding cell model created with stick filters from POm and VPM to three other models (Fig. 6E): i) with only stick filters from POm (n=7), ii) with sticks and sweep filters from POm and VPM, (n=22), and iii) with only sweep filters from POm and VPM (n=5). We observe that the model sweep-responding cell filters in layer 5a created with filters only from POm have a significantly larger error compared to the sweep-responding cells created with filters from POm and VPM (paired-t test, sticks, p<0.05, sweeps, p<0.01 or sticks + sweeps, p<0.02). It means that sweep-responding cells integrate stick-responding cells information from multiple structures. We also see that models created only with sweep filters or with stick filters and sweeps are not significantly better than models with only stick filters. This shows that stick information from the two thalamic regions is sufficient for creating sweep-like information in layer 5a of the cortex.

### Sweep coding in higher order POm is inherited from layer 5b and parallels layer 5a

It has been shown that layer 5a, where the new cortical sweep code emerges, is strongly innervated by higher order POm (10). We therefore recorded in this area, located 0.5 mm caudal to lower order POm (which we have called simply POm until now), to assess its contribution to the layer 5a sweep code. Repeating the correlated RC-VEL_W_ analysis used for lower order POm, we found a two-dimensional PC subspace accounting for around 80% of the filter variance (Fig. 7a). Remarkably, when we rotated and aligned the cortical subspace to the HO POm filter subspace, we obtained a quarter-turn rotation (44 degrees, Fig. 7b), meaning the HO POm feature space is mainly aligned with the sweep axis. In other words, HO POm contains only sweep-responding cells. Testing for directional tuning in this space (after rotating HO POm to match the cortical subspace and repeating the test of Fig. 5e), we again found rostral tuning for the sweep feature class (Fig. 7c, directional shuffling test, p<0.02). Studying the relative delay between matched filter subspaces across cortical layers, we found, to our surprise, that HO POm is only 0.9 ms in advance of layer 5a. By contrast, a 4.1 ms delay separates layer 5b from HO POm, making it plausible that the sweep code is inherited in part from layer 5b. To test this, we modeled HO POm sweep-responding cells as we did for layer 5a, here using the layer 5b sweep-responding cells we recorded as input. The model reproduced the sweep filter shapes well, with a delay of (−1.88±1.88ms), as expected, and a sigma distribution of (4.94±3.86ms). The alpha and error distributions are shown in Fig. 7f. The error distribution in this model has larger values than for the modeled layer 5a cells, suggesting, as we hypothesize, that sweep responses in HO POm likely draw on inputs beyond layer 5b.

**Figure 7.**
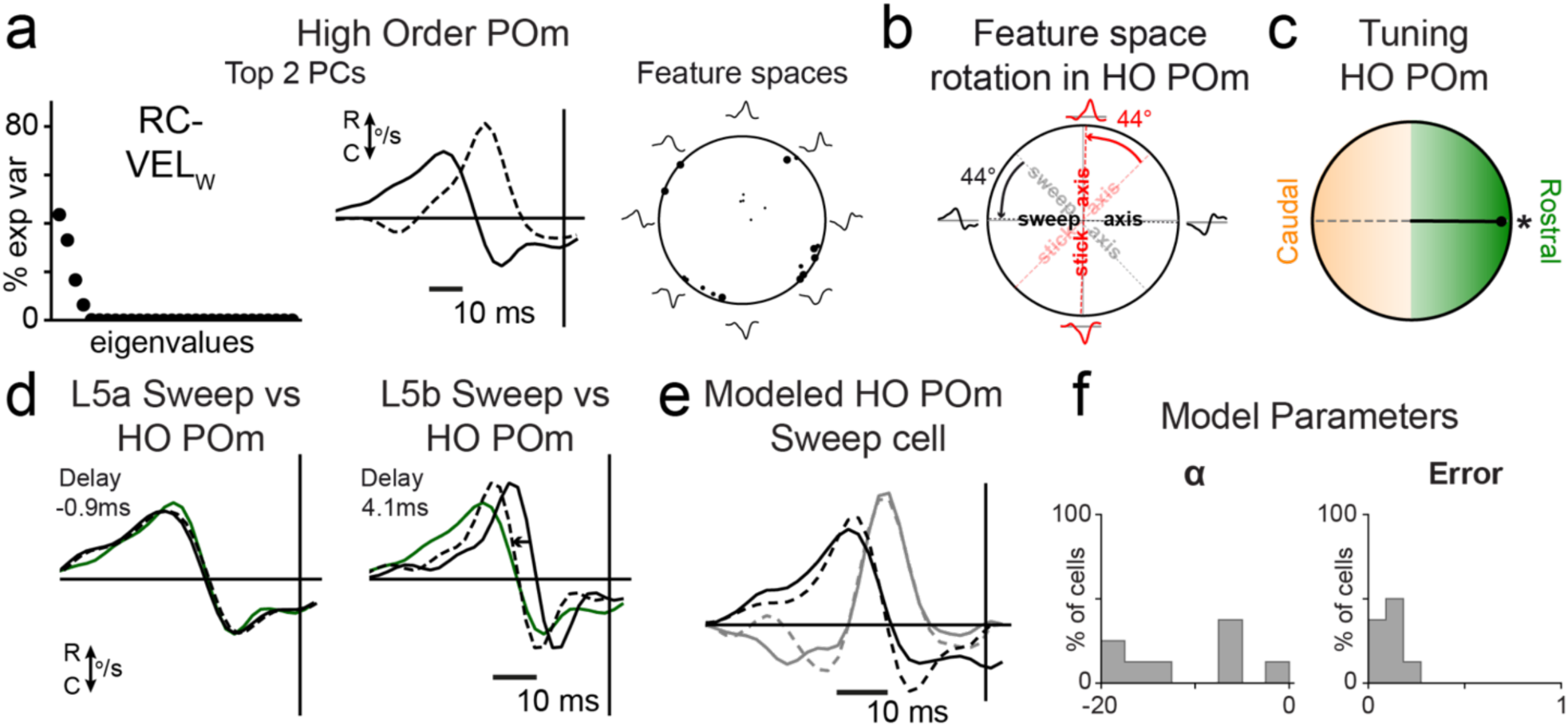
Higher-order POm (HO POm) carries a sweep code inherited from layer 5b and parallel to layer 5a. **(A)** Response subspaces obtained by principal component analysis made on the filters of Higher Order POm (n=23). Left: Variance explained. Center: First (bold) and second (dashed) PCA components. Right: Projection of all significant filters from all recorded cells onto the two-dimensional subspace defined by the population PCA. **(B)** Stick-Sweep feature space in HO POm, obtained by a 44° rotation of wS1 feature space. **(C)** Tuning of HO POm sweep- responding cells. Line represents the vector sum of all tuned cells. HO POm is significantly tuned rostrally (directional shuffling test, p<0.02). **(D)** Matching by rotating and delaying one PCA feature of HO POm (n=23) to the sweep features of specific layers in wS1 (L5a: 80 cells, L5b: 51 cells). **(E)** Model of one sweep-responding cell of HO POm created with neurons from layer 5b. (1st filters: black, 2nd filters: gray, model: dashed lines, experimental: solid lines). **(F)** Histograms of the fitted parameters for the HO POm sweep cell model: sweep-responding cells of HO POm (n=8) modeled by sweep-responding cells of layer 5b (n=12).

## Discussion

To sense the world, mice use their whiskers in an active process. This allows them to localize objects in space and to recognize shape and texture (6). How the brain integrates these tactile features remains a central question. Our results reveal that two distinct types of whisker movement features, sweeps and sticks, are represented in specific cortical layers of wS1. Sticks correspond to sharp, mono-directional deflections of single-whiskers, while sweeps are broader multi-whisker bidirectional movements extended over time (Fig. 3). Computationally, these correspond to two distinct temporal filtering operations: stick-responding cells act as fast, temporally precise detectors of brief single-whisker events, whereas sweep-responding cells act as temporal integrators that combine multi-whisker input over an extended window. Stick responses are inherited from VPM and predominantly represented in layers 3 and 4 of wS1 (Figure 4). In contrast, sweep responses emerge in layer 5a of wS1 through a broad temporal integration of stick-like signals from both VPM and POm (Figure 6). This sweep code is in turn relayed to higher order POm, which carries sweep responses with a latency similar to layer 5a, yet inherits them at least partly from layer 5b (Figure 7). These findings support the idea that laminar specialization in wS1 underlies distinct roles in tactile feature processing (23).

These features can be understood in the context of natural whisker-based exploration. When sensing an object, multiple whiskers are deflected, triggering stick-like responses that are encoded in VPM and relayed to layer 4 with a caudal directional bias (Fig. 5C). This supports the existence of an early object detection mechanism which operates both in thalamus and cortex (1). The anticorrelated velocity tuning between principal and secondary whiskers in stick-responding cells (Fig. 5 and Supplementary Fig. S7c) could further sharpen this detection, helping localize an edge or object more precisely at first contact. Afterwards, when the whiskers get released from the object, the motion transitions into bidirectional sweep-like movements, which are encoded in layer 5a in wS1, with a rostral preference (Fig. 5E). Sweep feature detection thus reflects a more integrative computation, likely suited for texture recognition, requiring prolonged temporal integration across multiple whiskers. On the contrary, stick coding, essential for rapid detection, is a process that relies on a smaller time window (Fig. 6D).

The whisker system relies heavily on bottom-up sensory pathways, particularly under our experimental conditions involving light anesthesia. This choice of preparation is dictated by our central aim, the identification of the precise velocity-defined stimulus features driving thalamic and cortical neurons. Reverse correlation to a high-dimensional, multi-whisker white-noise stimulus requires long, stable recordings and a fixed, exactly known mapping between the imposed stimulus and the neuronal response. Self-generated whisking would continuously alter whisker kinematics and contact conditions, confounding this mapping, which is why light anesthesia, as in foundational feature-selectivity studies of this system (13, 15, 17, 33), provides the controlled conditions the approach requires. A limitation of this design is that it does not capture how these features are exploited during active sensing, which awake behaving experiments will be needed to address. At the same time, top-down modulation is needed to regulate it: whisking is an active sensorimotor behavior that involves cortico-thalamic loops. wS1 integrates both externally driven touch and internally generated whisking signals, including self whisker deflections responses (24), as well as whisking phase-locked cortical responses (25). POm receives motor-related input and strongly innervates layer 5a of wS1 (Fig. 6A) (10), placing it in a strategic position to modulate sweep encoding. Although the precise role of higher order POm is still to be determined, several studies suggest a complex role in both amplifying and suppressing sensory signals. Inhibition of the cortex abolished spiking in POm from whisker movement (26), and POm activity plays a role in inhibiting layer 5a (27) and can also enhance the sensory signals (28). Notably, it was shown that POm activity is contingent on cortical input, as it has strong innervation from layer 5b of wS1 (29) and, as shown recently in awake mice, its responses to passive touch are markedly attenuated by barrel cortex inhibition (30). This is consistent with our finding that sweep responses in higher order POm can be partly reconstructed from layer 5b activity (Fig. 7). We hypothesize that the sweep encoding that emerges in layer 5a results from integrating both first-order stick information from first order POm and VPM, potentially combined with higher order contextual information. Future works aiming at understanding the features encoded in higher order POm with naturalistic stimuli could allow us to better elucidate the role of this structure.

Supporting this idea, recent work using optogenetic inhibition of higher order POm impaired texture discrimination in mice (31), consistent with our proposal that POm-related sweep integration underlies texture processing. This suggests that POm contributes to texture perception by integrating self-generated motor signals with tactile input. Previous studies have shown that thalamic spike timing encodes vibrissae dynamics with high temporal precision using only a few spikes (13). In our data, we show that VPM encodes fast stick-like movements with precise temporal profiles (Figure 4), whereas sweep-like integration of distributed inputs happens predominantly at the cortical level at longer latencies.

Our results can be placed within the longstanding model of the whisker thalamocortical system, in which the first-order VPM (lemniscal) pathway is associated with single-whisker coding, while the higher-order POm (paralemniscal) pathway is associated multi-whisker responsess (5, 7, 13). Our data agree with the first part of this model: VPM encodes fast, single-whisker stick-like movements with precise temporal filters, and these responses are relayed to layer 4 (Figure 4 and Figure 6). However, our findings refine the role attributed to POm, besides its previously described response to multiple whiskers. Rather than carrying prolonged, multi-whisker sweep responses directly, first-order POm predominantly contained stick-responding cells, much like VPM. The multi-whisker, temporally extended sweep code does not appear to be inherited ready-made from the paralemniscal thalamus, but instead emerges within the cortex, in layer 5a, through temporal integration of stick-like inputs from both VPM and first-order POm (Figure 6). Higher-order POm does carry sweep responses, but with a latency similar to layer 5a and inherited at least in part from layer 5b (Figure 7), placing it downstream of the cortical computation rather than as its source. Thus, our results position the emergence of multi-whisker sweep coding from the thalamus to the cortex, while preserving the classic view of VPM as a fast, single-whisker channel.Our findings also relate to prior work in wS2, which exhibited longer temporal integration compared to wS1 (18), possibly supporting the computation of global motion across many whiskers. However, the specific movement features could not be resolved in that study. Applying our velocity-based stimuli to study the responses of wS2 could help test whether sweep-like responses are even more prominent in this area.

A major methodological contribution of our study is the design and comparison of multiple stimulation protocols to identify the most selective one for extracting response features. We found that Gaussian rostro-caudal white noise in velocity (RC-VEL_W_) is the stimulus that elicits the most selectiveness (Fig. 2). Importantly, this stimulus did not increase overall firing rate compared to others, such as white noise in acceleration, but instead produced more informative stimulus subspaces. Concerning the white noise in position stimulus, it produced drastically fewer filter features and fewer spikes than a previous study in rats, likely due to the smaller angle space explored by this stimulus, but also due to differences in whisker diameter and torque. The torque exerted on mice whiskers may be significantly smaller compared to the one in rats when using the piezoelectric Matrix (32). The POS2_W_ stimulus has lifted this problem by eliciting more spikes and producing more filters, but lacked selectivity due to its narrow acceleration range, producing highly correlated stimuli across time (Fig 2E). This further highlights the advantages of the RC-VEL_W_ stimulus in resolving fine temporal structure.

Directionality also plays an important role. Previous studies have found that wS1 exhibits a direction preference for global motions of whiskers in the cortex and VPM (33,34), and that the cortical column seems to be more tuned to the caudo-ventral direction (35). Here we show that the most selective stimulus is a rostro-caudal one, especially along one direction (either rostral or caudal). VD-VEL_W_ also shows high selectiveness, though in a smaller degree (Supplementary Fig. S2). The cortex encodes with better detail rostro-caudal movements, in close relation with the rostro-caudal direction of whisking movements.

Texture recognition is a demanding computational task requiring integration across both space and time, and mice perform well at recognizing textures (36,37). How the texture is encoded in the brain can help us better understand how the whisker system is integrating information. It is now well established that the cortex is needed to decode texture (38,39), and that POm, and particularly its interactions with wS1, is also essential for this function (29). Therefore, these studies support the existence of a sensorimotor loop involving higher order POm and the layer 5a and 5b of wS1 for being able to discriminate textures (40), consistent with the sweep code we find shared between layer 5a and higher-order POm (Fig. 7). We propose that the sweep-responding cells in layer 5a may reflect the integration of current sensory input with motor-related signals from higher order POm. This could help synchronize texture processing to moments of actual contact, rather than during free-air whisking. One remaining question is how and when this computation is initiated. We speculate that the motor cortex could guide this computation by sending timing cues, when an object is encountered, to POm, which in turn modulates cortical sweep-responding cells. It is known that wM1 contributes to object localization by encoding the position of the whiskers (6), and wS1 actively terminates whisking and provokes the retraction of the whiskers, upon object contact. These observations further support the need for a closed sensorimotor loop linking motor cortex, thalamus and sensory cortex to guide active tactile perception.

Our findings show that different layers of the barrel cortex encode distinct whisker features: fast, directional stick coding dominates in layer 3 and 4, while slow, multi-whisker sweep coding emerges in layer 5a through temporal integration of thalamic input. This layered organization suggests a division of labor between rapid detection and integrative processing. Sweep-responding cells may support texture discrimination by combining sensory and motor-related signals, especially via higher-order thalamic input from POm. While our recordings targeted the VPM and POm relay nuclei that provide feedforward input to the cortical layers studied here, dissecting the broader thalamocortical loop and its corticothalamic contributions, including through direct manipulation of these pathways, remains an important direction for future work. Future studies in awake, behaving animals will be key to understanding how these features contribute to sensorimotor computations during natural whisking.

## Materials and Methods

### Animal procedures

Mice experiments were in conformity with the French Ethical Committee and European legislation. 12 mice were used (5 for the thalamus recordings, 7 for the cortex) aging 4 to 6 weeks, weighing 25.85 ± 1.42g. Anesthesia was induced with 3% of Isoflurane in O_2_, and then the animal was head fixed on a stereotactic device. The body temperature was maintained at 37°C with a controlled heating pad and a rectal probe. To prevent drying of the eyes, Ophtalon (optical gel) was applied. The whiskers were cut at 5mm long and then connected to the Matrix stimulator. During the experiment, the anesthetic state was monitored with ECoG electrodes to keep it in a light regime. These electrodes were inserted through two small craniotomies (0,5mm), placed anterior to the multi-electrode recording site.

For the barrel cortex experiments, we first performed intrinsic imaging to localize the barrel C2. Then, a craniotomy (2mm diameter) was performed centered on the C2 signal (Supplementary Fig. 1D) position (near 3,3mm lateral –1,5mm posterior from Bregma) and the dura was removed. Extra care was taken to prevent swelling on the durectomy site by minimizing its size as much as possible. Multi contact silicon probes (64 channels Neuronexus tetrode 4x4 or Buszaki 64 with 4 shanks) were lowered to reach the different layers of the barrel cortex (150 μm – 1300 μm) at a slow pace (<1μm/s).

For the experiments in the lower order thalamus, a craniotomy (2mm diameter) was done on the left hemisphere (3,5mm lateral –1,5mm posterior from Bregma) and the dura was removed. A positioning at 2 mm posterior from Bregma was used to reach higer order POm. The same type of silicon probes used in cortex were lowered to reach VPM and POm. During the slow lowering procedure (<1μm/s), spike signals recorded by each electrode were connected to an audio device and a correlated periodic sparse movement was applied to the whisker pad. This allowed us to know when the electrodes reached the thalamus as a rhythmic sound was immediately recognizable (Supplementary Fig. 1F)

The isoflurane concentration was decreased gradually to 1-1.2% to change the anesthetic state to light (stage III, plane 1–2) when the lowering of the electrodes was finished. The anesthetic state of the animal was monitored with the ECoG signal (> 5 Hz), the respiration rate (between 1-1.5 Hz) and the lack of spontaneous movements.

### Intrinsic Imaging

For cortical measurements, intrinsic images (taken at a frequency of 20Hz) were obtained using light of 630 nm and a coupling of a Qicam CCD camera (Q-imaging) with a Leica MZ9.5 microscope (Supplementary Fig. 1C-D). Piezoelectric stimuli of the C2 whisker were applied at 10 Hz for 1s during the recording. The intrinsic images were obtained by subtracting the images just before the stimulus to the images during the stimulation. These images gave us the localization of the C2 barrel in the cortex and helped us to define the position of the craniotomy and thus the entry point of our probe.

### Electrophysiological recordings

During the experiments, 64 raw electrophysiological traces (2.5 h per recording) were recorded at 30kHz with a Blackrock amplifier. The pre-processing of the data was done with the Klusta suite (41). With this software, spikes were automatically extracted and sorted thanks to their waveforms. During offline analysis, the phy viewer suite was used to do a manual curation of the data. Each group of spikes was viewed and categorized as single units, multi-units or noise. A group of spikes composed of non-neuronal activity was categorized as noise. A group of spikes that contains multiple waveforms that could not be separated was categorized as multi-units. A group of spikes well defined in space, in shape, with a minimum number of spikes (>400) and with an auto-correlogram showing a refractory period was defined as deriving from a single unit. Only the single units were used in the following results. For the recordings in the cortex, we obtained 405 single units from 7 experiments. For the recordings in thalamus, we obtained 453 single units from 5 experiments. We aimed the different recordings across animals specifically to the middle of the cortical layers, which produced a lower density of recorded cells in the limits between layers (Fig. 3E).

### Stimulus application

A piezo-electric based stimulator was used to deflect the whiskers in a controlled way. The Matrix is a custom-built whisker stimulator and was developed in the laboratory (Supplementary Fig. 1A-B) (14). The whisker trajectory is controlled vertically and horizontally with millisecond and micrometer precision. The controlled deflection of the whiskers can be done with a frequency up to 1kHz. The resonance vibrations (ringing) caused by the high frequency movement is compensated numerically with custom made software for the Matrix. Nonetheless, the ringing and the physical constraints compelled us to create a stimulus with a frequency cut-off of 80Hz. During the experiment, after the mice were anesthetized and positioned, the 24 caudal whiskers were cut at 5mm and inserted inside each piezo-electric deflector of the matrix. The deflectors were positioned to have the tip at 2.5mm of the skin and aligned to the resting position of the whiskers. The resting position was defined as the zero position for our stimulus.

For the experiments in the thalamus and in the cortex, the following stimuli were applied. First, sparse noise in position (deflections of one whisker at a time) was applied vertically and horizontally (cortex). In the thalamus, sparse noise in velocity (deflections in velocity of one whisker at a time) was applied after the sparse noise in position. Second, we applied different types of Gaussian white noise in periods of 10 seconds. Each type was applied 200 times in a randomized order. For the cortex, we had two batches of experiments with different stimuli. In the first batch of 4 experiments, we applied RC-VEL_W_, POS2_W_ and ACC_W_. In the second batch of 3 experiments, we applied RC-VEL_W_, POS_W,_ VD-VEL_W_ and uncorrelated RC-VEL_W_. For the thalamus experiments, we applied RC-VEL_W_, POS_W_ and POS2_W_ (Supplementary Fig. 1G). For the POS2_W_ stimulus, fewer spikes are needed to have significant filters, because this stimulus has a higher correlation across time. This higher correlation creates more defined movements that, when applying STA or STCs, will find well defined filters more easily.

To create Gaussian White Noise stimuli, we based ourselves on previous work (15). To make the different stimuli comparable, the probability distributions across them were equalized in velocity space, to explore the same values (Fig. 1B). Their white domain spectral content shows a flat power density up to 100 Hz (40Hz for the POS2_W_ stimulus).

### Histology

Before the lowering of the electrodes, DiI was applied on the electrodes to allow visualizing the trajectory of the electrodes in the brain at the end of the experiment. After recordings were ended, a pentobarbital overdose was injected to euthanize the animal. Then, a transcardiac perfusion of saline and formaldehyde were performed to fix and preserve the brain tissue.

For the cortex experiments and to localize the electrode position and determine its depth, multiple tangential slices of 100 µm were taken. The depth of recording for each electrode contact was computed with the reading of the lowering of the probe and by taking into account a 5° deviation from the vertical orientation.

For the thalamus experiments, 80μm thick coronal brain slices were cut and stained with cytochrome oxidase to localize the path of the electrodes, and then the position of the recorded cells. The goal was to include in our analysis only the cells that were in VPM or in POm. The coronal slice with the most visible probe shanks was taken and matched with the Allen brain Atlas (Dong et al., 2008) to locate POm and VPM. Each single electrode was positioned along the trajectory of the shanks thanks to the known depth and the geometry of the electrode. The position of a neuron was defined as the position of the contact where the maximum waveform was found. We recovered 105 neurons in first order POm, 43 neurons in VPM and 23 neurons in Higher Order Pom out of the 453 isolated single units.

### Reverse correlation analysis

All the analysis were performed offline and coded in the Python language. Windows of 40 or 60 ms (40 ms for thalamus, 60 ms for cortex) of whisker stimulus were taken around each spike (35 or 55 ms pre- and 5 ms post-spike). A spike trigger ensemble is therefore created for each cell containing all these windows of stimulus, on the kinematic feature corresponding to the white stimulus domain (position for POS, velocity for VEL, acceleration for ACC). If a neuron is encoding a particular shape of whisker deflections, we can assume that this shape should cause its firing more often than by chance. Whitening was applied to each whisker deflection of this spike triggered ensemble by multiplying each whisker deflection by a whitening matrix calculated for each stimulus. Averaging the spike triggered ensemble gives us the spike triggered average (STA). Principal Component Analysis was applied to the spike trigger ensemble to extract the multiple shapes of whisker deflections that explain the most the variance of the entire ensemble, a procedure called Spike Triggered Covariance (STC). To assess the significance of these shapes, the same analysis was repeated 200 times with random shuffles of the stimulus and the same spike trains. The whisker deflection shapes were categorized as significant if they had an eigenvalue above seven standard deviations of the 200 random shuffling average. A cell can have significant STCs without having significant STAs. As an example, if significant filters out of an STC produce spikes in a cell for both directions (rostral and caudal for example), therefore, an average of these filters can create a non-significant null STA.

### Feature spaces and functional selectivity

To characterize the feature space shared across the recorded population, we performed a second principal component analysis, this time on the significant filters pooled across all neurons, rather than within a single cell as in the per-cell STC described above (as was done previously in (15), (17) and (18)).. For each stimulus type, two population filters (PCs) were significant and explained 90% of the variance. With this encoding, each cell’s significant filter can be mostly explained as a linear sum of these two PCs. Therefore, we created polar feature spaces out of these two filters. In this feature space, each significant filter of a cell can be represented as a point (the point size is proportional to the significance of each filter for the cell).

For each cell, we looked at their individual tuning in this feature space. A polar-radial histogram of firing rate was done by calculating the number of spikes falling in each portion of the polar-radial binning. Each spike has an angle and radius, and it can be considered as a vector in this representation. To assess the significance of this tuning, we tested it by shuffling 1000 times the angles of the spikes while conserving their radial values. To be called region specific, a vector sum of the tuning of a cell needed to have a radius above 95% of the random shuffles. We also tested bidirectional selectivity by only using half of the circle by multiplying every angle by two and by performing the same significance test.

### Sparse noise analysis

The sparse noise stimuli were used for multiple purposes in our analysis. First, it allows us to obtain the receptive fields of each cell, determining a principal whisker and surround whiskers. This is calculated by computing a peristimulus-histogram for each whisker and for each direction used. Second, we use the sparse noise to classify between stick and sweep-responding classes. A cell was deemed as a stick-responding cell if for one of the whiskers, the number of spikes occurring for 15 ms after the onset of the stimulus was 5 times the baseline rate. The baseline rate was extracted for blank periods where no stimulations were present. To classify a cell as sweep-responding, the Z-score mean firing rates were computed for the 60 ms following every whisker deflection. The mean and standard deviation used was extracted from the blank period. The Z-score above 0.5 was summed and multiplied by a sharing parameter defined in (15). This number called RF was then compared to 1000 shuffles of the same RF calculated with the basal rate of the whisker without deflection. If the RF calculated was above every shuffled RF, the cell was defined as a sweep-responding cell.

### Matching STC across structures

We matched STC filters, stick or sweep curves between structures to find if there were similarities between them. To do this, we fixed the curves in the cortex and we changed the curves in the thalamus by rotating them in the feature space and by delaying them in time. The goal was to find the best match, which was obtained by maximizing the energy of the sum of the two scalar products between the rotated and delayed filters of the thalamus and the fixed curves of the cortex. With this method, we could find the best match for each curve (STC, stick or sweep filters) and the optimal delay and rotation.

### Waveform classification

To classify cells as fast and regular spiking in the cortex, we extracted 7 features for each waveform of each neuron (Supplementary Fig. S6A). The waveform used for each cell was the mean across spikes in the electrode where we recorded the maximum signal amplitude. A principal component analysis was applied on these features for each neuron. In this space, the first three PCs can explain 85% of the variance (Supplementary Fig. S6B). We plot each neuron as a point in the space created by the two first PCs for a 2D visualization. We applied the k-nearest neighbor method to find 16 groups in this space. After this automatic overclustering to guarantee no cross contamination between groups, we assigned each group into three classes (fast, regular or undefined) using the 2D visualization and by coloring each cluster. Finally, to verify this classification, we plotted each group in the feature space and their waveforms (Supplementary Fig. S6D). The neurons in the thalamus are reported to be predominantly regular spiking. To verify this, we did another principal component analysis including both the waveforms of the neurons in the cortex and in the thalamus. We obtained a new feature space and we color plotted the already done classification of fast and regular spiking cells in the cortex. We plotted, together with cortical neurons, the neurons of POm (low and high order) and VPM (Supplementary Fig. S6E, here visualized with a PC1 and PC3 projection). We see that the neurons in the thalamus are mostly regular spiking as expected.

### Rostral/Caudal tuning with Gaussian white and sparse noise

In the feature space obtained by the RC-VEL_W_ stimulus, we can define the rostral and caudal axis based on the quota between the area above and below the curves. To assess the rostro-caudal tuning of regions, we sum the normalized tuning of each cell in the feature space obtained by the RC-VEL_W_ stimulus. To know if this rostro-caudal tuning is significant, we do 1000 shuffles of random angles sums (same number as cells in the studied structure) and compare to the original radial value The cell population is tuned if its radial value is above 95% of the shuffles populations to be deemed significant.

To test the rostro-caudal tuning, we also used sparse noise to define a velocity rostro-caudal ratio. This ratio was defined as the number of spikes responses to positive velocity (first 20 ms of the sparse noise whisker deflection) subtracted by the number of spikes during negative velocity (the following 20 ms). This ratio was then normalized by the sum of all spikes (40 ms). This rostro-caudal ratio was obtained on all whiskers for each cell. Afterwards, we can look at the distribution of this ratio for each structure. We used a Wilcoxon biased test for significance.

### Model of Stick and Sweep cells

To model the responses of stick-and sweep-responding cells, the first two filters of the spike-triggered covariance (STC) responses to the RC-VEL_W_ stimulus (white noise in velocity) were reproduced (Fig. 6C and Fig. 7E show three example fitted cells). A model of layer 4 stick-responding cells was created from VPM stick-responding cells, and four different models of layer 5a sweep-responding cells were created from POm/VPM stick- and sweep-responding cells, and one model of higher-order POm was created from layer 5b sweep-responding cells. Up to two significant filters of the STC response to the stimulus were used for each modeled cell (for example, to model layer 4 stick-responding cells, only the first filter of VPM stick-responding cells was used). These filters were delayed randomly following a skewed Gaussian distribution of delays. A binomial law with a probability of 0.95 was then applied to randomly change the sign of some filters. These delayed filters formed a spike-triggered ensemble (Fig. 6B). We used between 750 and 1700 delayed filters in the ensemble, comparable to the number of filters in the spike-triggered ensemble of real cells. Finally, a principal component analysis was computed to obtain the modeled response of the cell.

The three parameters of the skewed Gaussian distribution (mean, standard deviation, and skewness) were explored to fit the STC responses of each cell. The mean and the skewdness was in the range of [-20,0] with steps of 0,5. The standard deviation was in the range of [0,15] with steps of 0,5. Delays were constrained between −30 and +10 ms. The error was defined as one minus the sum of the scalar product of each modeled filter with the corresponding real-cell filter, weighted by the normalized eigenvalues of the real cell (to better fit the filters explaining the most variance). This error is plotted in Fig. 6D and Fig. 7F. For some models, in order to reduce the error, and emphasize part of the filters that were poorly fitted, we added a weight of two over a specific portion of the filters for the error calculation as a constraint. For the layer 5a sweep model, a portion of the first filter between −5 and +5 ms was weighted; for the HO POm sweep model, a portion of the two filters between −15 and −5 ms was weighted.

### Uncorrelated Reverse correlation analysis and tuning

An uncorrelated RC-VEL_W_ stimulus was applied to the whisker in 3 animals, from which we recorded 254 cells. Each whisker is stimulated by a RC-VEL_W_ stimulus that is uncorrelated from the stimulus applied to the other whiskers. Therefore the same analysis used for analysing the RC-VEL_W_ was applied for each whisker. A principal component analysis was applied to extract the deflection shapes explaining most of the variance of the entire spike-triggered ensemble, for each whisker (Supplementary Fig. 7a, b, left). For this stimulus, two population filters (PCs) were significant and explained almost 90% of the variance. With this encoding, each cell’s significant filter can be mostly explained as a linear sum of these two PCs. Therefore, for each cell and each whisker, we looked at their individual tuning in this feature space (SupFig 7a, b, right). To be considered significant, a whisker’s tuning needed a radius above 95% of the 1000 random shuffles.

To look at the tuning angle distribution, the difference between the Principal whisker tuning angle and the secondary whisker tuning angle were computed for each cell. The principal whisker was defined as the significant tuning with the highest radius. The secondary whisker was defined as the neighboring whisker with the highest radius. The difference in tuning angle was computed and plotted for all cells, stick-, sweep- and mixed responding cells (Supplementary Fig. 7F).

## Acknowledgments

The authors thank Guillaume Hucher for performing the histology and Aurélie Daret for providing the animal care.

## Author contributions

Conceptualization: DES, MAG Methodology: TQ, MAG Investigation: TQ, MAG Visualization: TQ, MAG Supervision: DES, MAG Writing—original draft: TQ, MAG

Writing—review & editing: TQ, DES, MAG

---

**Supplementary Figure S1.**
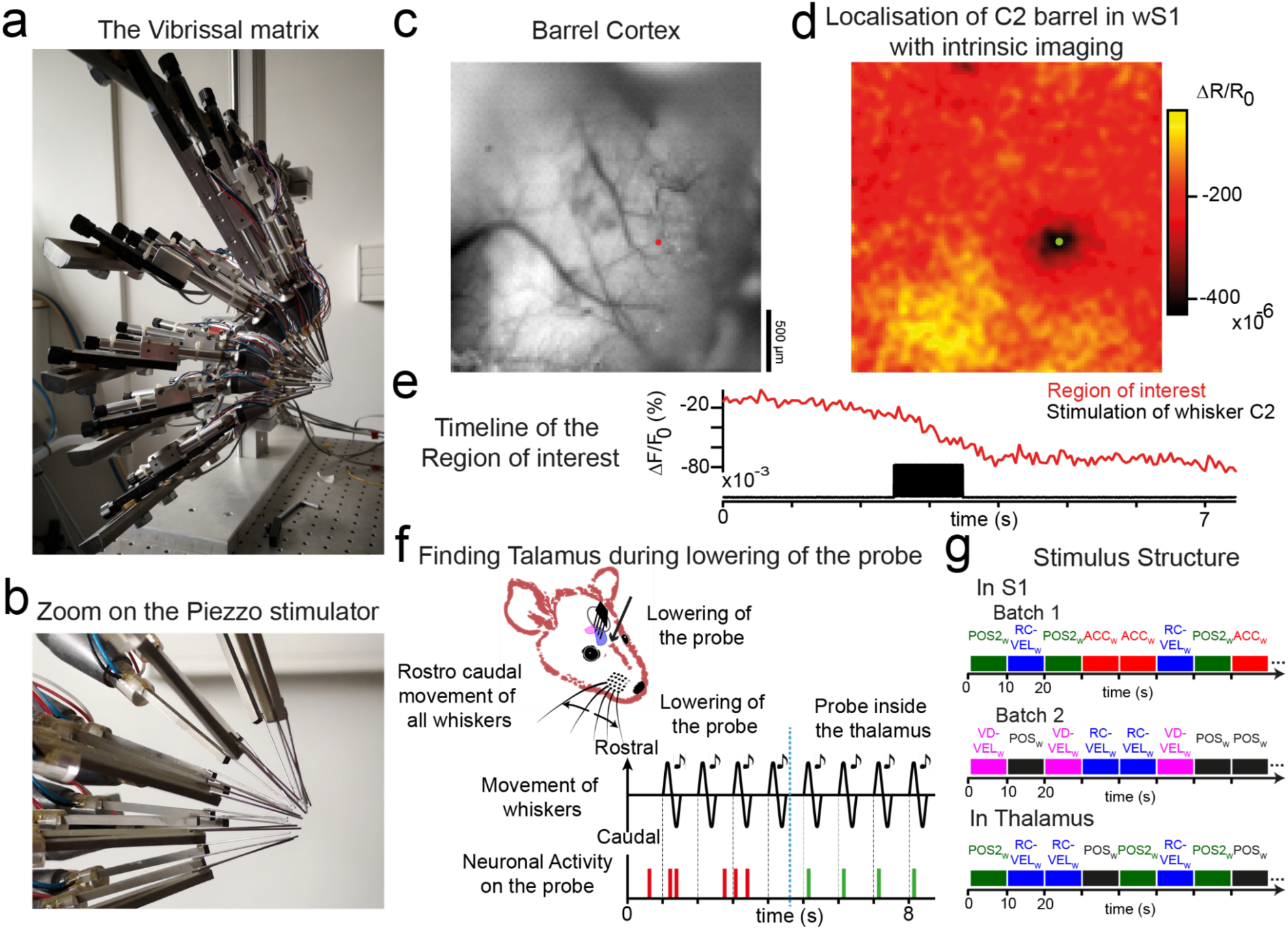
Experimental details. **(A)** Photo of the matrix of piezo-electric stimulators used for the deflection of the whiskers. **(B)** Zoom on the piezo-electric stimulators. **(C)** Photo of the barrel cortex before intrinsic imaging. **(D)** Results of the intrinsic imaging technique used to locate the C2 barrel in wS1. **(E)** Timeline of the response in the region of interest (C2 Barrel in wS1). **(F)** Description of the method used to find VPM and POm in the thalamus. During the lowering of the probe, all the whiskers were stimulated periodically with the same movement. A sound was played at the onset of movement. By looking at the neuronal activity on the probe, we could find a spot where the activity was synchronized with the sound, and therefore the whiskers movements. **(G)** For the cortex experiments, two batches of stimulus were applied, these batches were composed of multiple 10 seconds Gaussian white noise stimuli in a randomized order. For the thalamus experiments, only one batch of stimuli was played with the same structure.

**Supplementary Figure S2.**
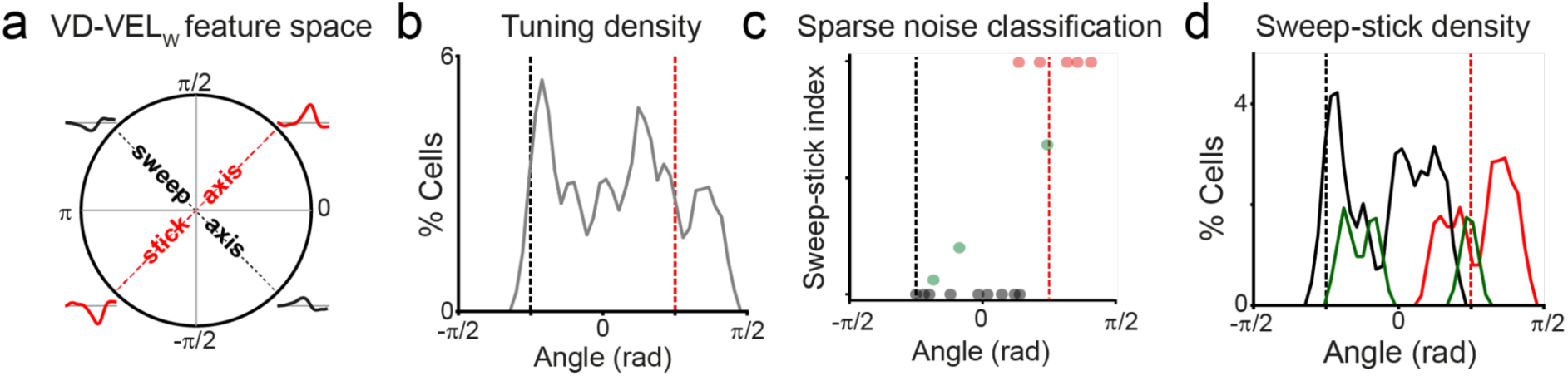
Classification of stick/sweep-responding cells in the velocity ventro-dorsal feature space. **(A)** The velocity ventro-dorsal features space (stick/sweep axis defined as in the rostro-caudal feature spaces) **(B)** The density of cells tuned to a specific angle in the Sweep-stick ventro-dorsal feature space **(C)** Classification of sweep (black), stick (red) and mixed (green) -responding cells using a Sparse Noise stimulation sweep-stick index. **(D)** Distribution of sweep (black) stick (red) and mixed (green) -responding cells across the Sweep-stick ventro-dorsal feature space.

**Supplementary Figure S3.**
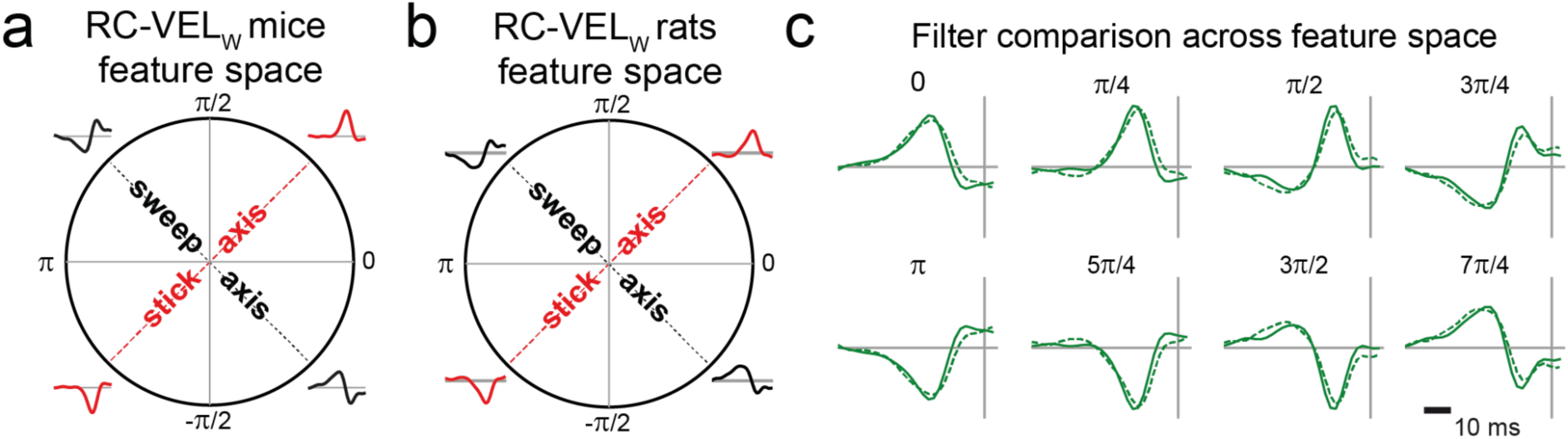
Comparison between rats and mice sweep/stick feature space. **(A)** Mice sweep/stick feature space **(B)** Rats sweep/stick feature space **(C)** Comparison of filters between mice and rats across the feature spaces on different angles.

**Supplementary Figure S4.**
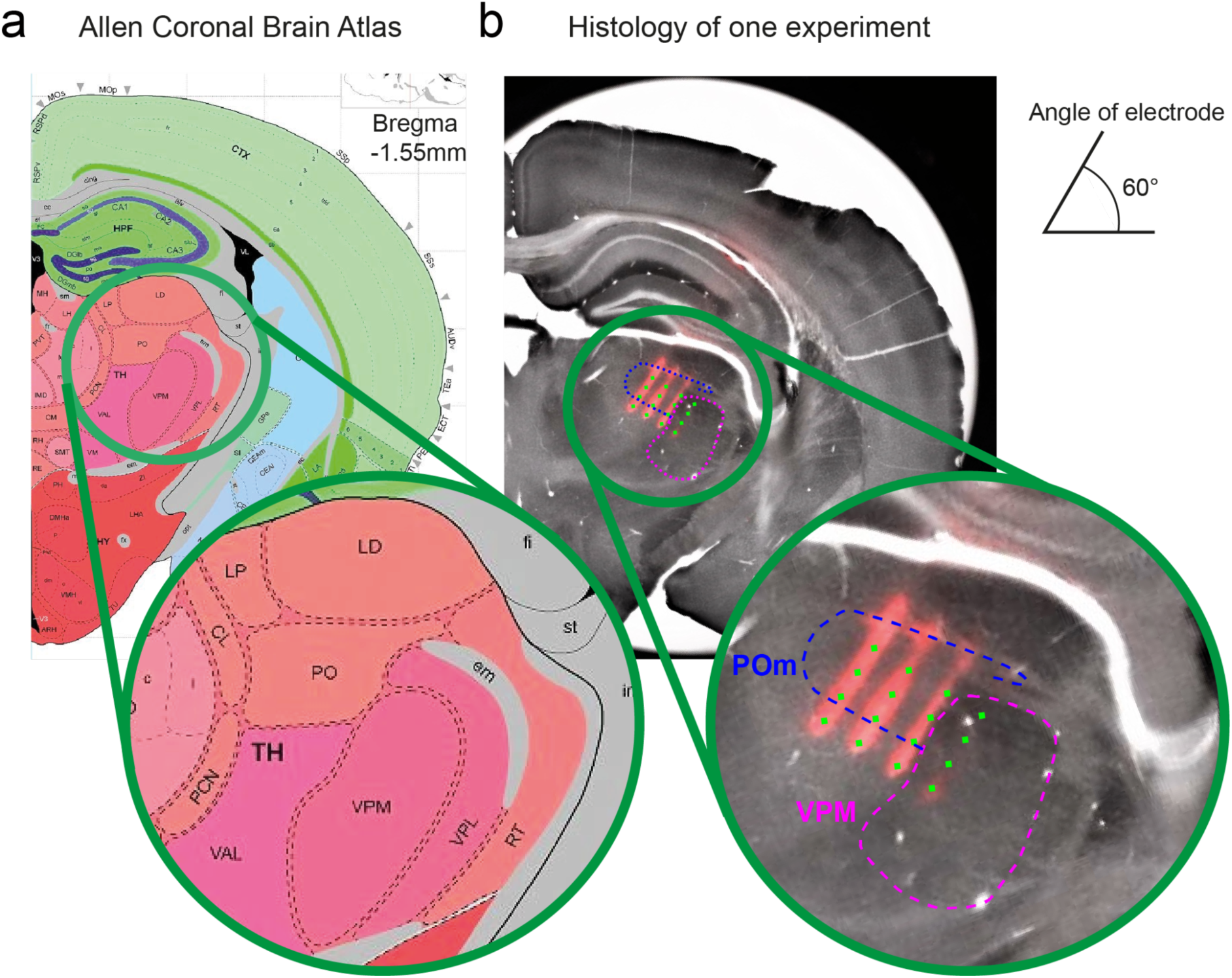
Histology analysis and comparison with an atlas. **(A)** Coronal slice on Allen brain atlas for the mouse brain (Bregma -1.55). **(B)** Slice of brain from a mousse where we see the end of the electrodes with DiI in red. On this slice, we visualize POm and VPM thanks to the corresponding slice in the atlas. By knowing the position of each shank of the electrode, we report their location on this slice (green).

**Supplementary Figure S5.**
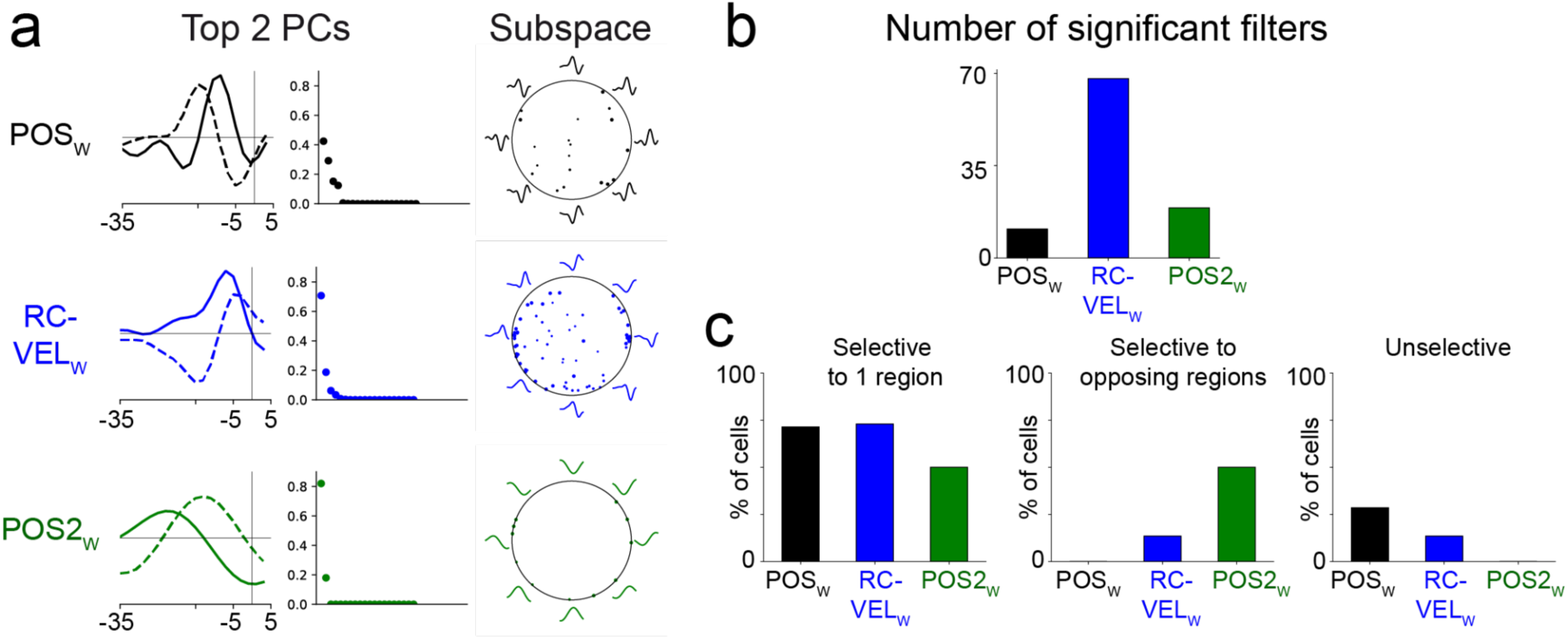
Selectiveness of different stimuli in the thalamus. **(A)** Top 2 filters and subspaces of different stimuli. **(B)** Number of significant filters for different stimuli. **(C)** Selectiveness of rostro-caudal velocity (blue), position (green), position2 (black) Gaussian white noise stimulus.

**Supplementary Figure S6.**
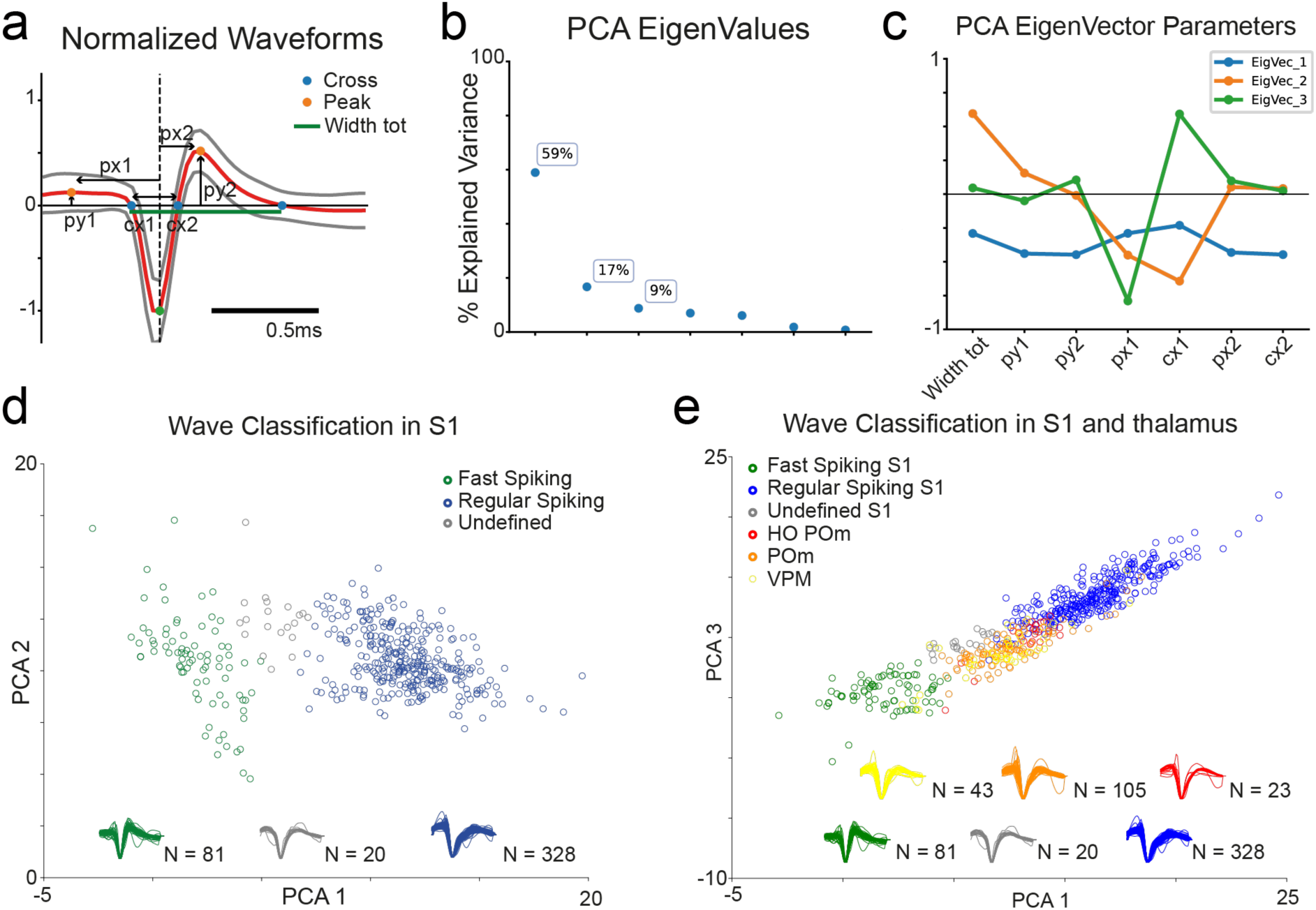
Spike waveform analysis (Fast and Regular Spike Classification) **(A)** Definition of parameters for the spike waveforms. **(B)** Eigenvalues of the different eigenvectors as a result of a principal component analysis. **(C)** Influence of each defined parameter for each eigenvector. **(D)** Spike-waveform classification in wS1 of fast (green), regular (blue) and undefined (grey) spiking cells. **(E)** Spike-waveform classification in wS1 of fast (green) and regular (blue) and undefined (grey) spiking cells. On top of it, spike-waveform in VPM (yellow) and POm (orange) of spiking cells.

**Supplementary Figure 7.**
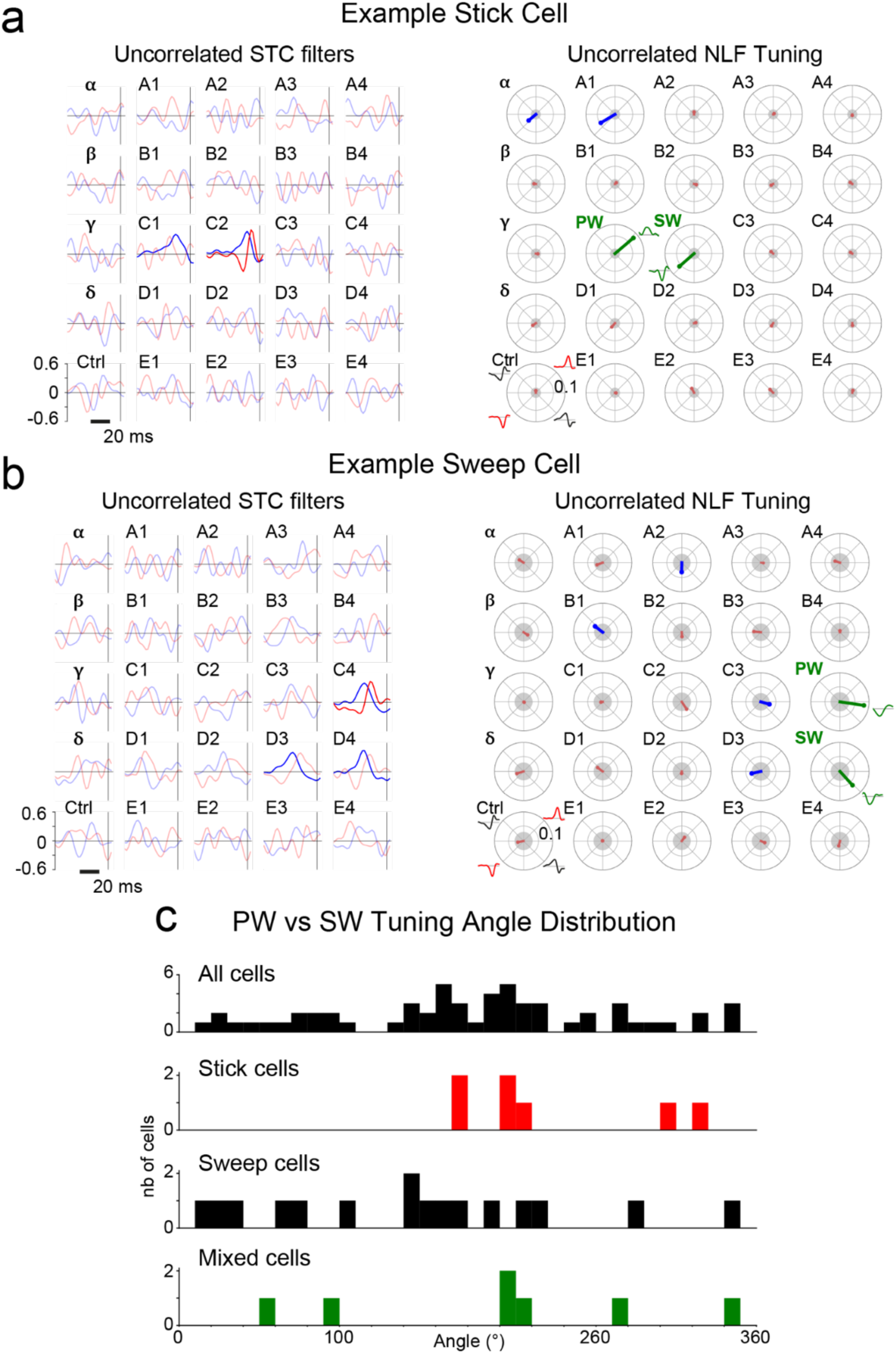
Stick-responding cells show anticorrelated principal/secondary whisker (PW/SW) tuning under uncorrelated RC-VELW stimulation. **(A-B)** Example STC filters and NLF tuning obtained from one stick and one sweep cell in response to uncorrelated RC- VELW. The control panels show the same analysis done on a stimulus not played to the whiskers. (Left) STC filters in response to uncorrelated RC-VELW. Significant filters are shown in bold (first filter: blue, second filter: red). (Right) NLF tuning in response to uncorrelated RC-VELW. Significant tunings are shown in blue (directional shuffling test, p<0.05). Insignificant tunings are shown in red (directional shuffling test, p>0.05). The two green tunings are the principal whisker and secondary whisker (PW and SW) which are significant with the highest radius (see Methods). They are used to calculate the tuning angle between PW and SW. **(C)** Histograms of tuning angle distribution between PW and SW for all cells (n=58), stick- (n=7), sweep- (n=16) and mixed- responding cells (n=7).

